# Stabilization of Interdomain Interactions in G protein α_i_ Subunits Determines Gα_i_ Subtype Signaling Specificity

**DOI:** 10.1101/2023.03.10.532072

**Authors:** Tyler J. Lefevre, Wenyuan Wei, Elizaveta Mukhaleva, Sai Pranathi Meda Venkata, Naincy R. Chandan, Saji Abraham, Yong Li, Carmen W. Dessauer, Nagarajan Vaidehi, Alan V. Smrcka

**Author notes:** Correspondence: Alan V. Smrcka, Ph.D. These authors contributed equally (Wenyuan Wei, Elizaveta Mukhaleva).

## Abstract

Highly homologous members of the Gα_i_ family, Gα_i1-3_, have distinct tissue distributions and physiological functions, yet the functional properties of these proteins with respect to GDP/GTP binding and regulation of adenylate cyclase are very similar. We recently identified PDZ-RhoGEF (PRG) as a novel Gα_i1_ effector, however, it is poorly activated by Gα_i2_. Here, in a proteomic proximity labeling screen we observed a strong preference for Gα_i1_ relative to Gα_i2_ with respect to engagement of a broad range of potential targets. We investigated the mechanistic basis for this selectivity using PRG as a representative target. Substitution of either the helical domain (HD) from Gα_i1_ into Gα_i2_ or substitution of a single amino acid, A230 in Gα_i2_ to the corresponding D in Gα_i1_, largely rescues PRG activation and interactions with other Gα_i_ targets. Molecular dynamics simulations combined with Bayesian network models revealed that in the GTP bound state, dynamic separation at the HD-Ras-like domain (RLD) interface is prevalent in Gα_i2_ relative to Gα_i1_ and that mutation of A230^s4h3.3^ to D in Gα_i2_ stabilizes HD-RLD interactions through formation of an ionic interaction with R145^HD.11^ in the HD. These interactions in turn modify the conformation of Switch III. These data support a model where D229^s4h3.3^ in Gα_i1_ interacts with R144^HD.11^ stabilizes a network of interactions between HD and RLD to promote protein target recognition. The corresponding A230 in Gα_i2_ is unable to form the “ionic lock” to stabilize this network leading to an overall lower efficacy with respect to target interactions. This study reveals distinct mechanistic properties that could underly differential biological and physiological consequences of activation of Gα_i1_ or Gα_i2_ by GPCRs.

## Introduction

Many physiologically important hormones and neurotransmitters signal through G protein-coupled receptors (GPCRs), rendering these membrane-spanning receptors highly clinically significant as important drug targets ^1, 2^. GPCRs transduce signals into the cell via heterotrimeric G proteins, consisting of the Gα subunit and the Gβγ constitutive heterodimer. Signaling diversity from GPCRs is primarily achieved via an array of Gα subunit protein families which harbor distinct downstream signaling capabilities, including the G_s_, G_i/o_, G_q/11_, and G_12/13_ families ^3–6^.

Gα subunits consist of a Ras-like domain (RLD), which binds and hydrolyzes guanine nucleotides, and an all-helical domain (HD), connected by a flexible hinge region ^5, 7^. Much of the investigative focus on Gα protein function has been on the RLD, which harbors three “Switch” regions (Switch I-III) that undergo conformational alterations upon GTP binding. Upon binding GTP, Switch regions I-III collapse toward the bound nucleotide in a conformational rearrangement that permits Gα·GTP-effector interaction after separation from Gβγ and the receptor ^8^. In contrast, the HD is relatively rigid and opens along the interdomain cleft via the flexible hinge in the nucleotide free transition state along the pathway of receptor-mediated GDP release ^9–11^. Mutation of residues along the Ras-HD interface further increases receptor-independent rate of GDP dissociation in Gα_i_^12^.

Generally, the Gα_s_ family activates adenylyl cyclases (ACs) to produce 3’,5’-cyclic adenosine monophosphate (cAMP) and the Gα_i_ family inhibits ACs ^3^. The Gα_i/o_ family consists of Gα_i1_, Gα_i2_, Gα_i3_, Gα_o_, Gα_T1_, Gα_T2_, Gα_T3_, and Gα_z_. Gα_o_ is prominent in the brain, Gα_T_ in the visual and taste systems, and Gα_z_ in the brain and prostate. Gα_i2_ protein expression is more widespread and more abundant than any other protein in the Gα_i/o_ family, except for Gα_o_^13^. Gα_i1-3_ are expressed broadly in humans, with Gα_i2_ often being expressed alongside Gα_i3_ and/or Gα_i1_. Gα_i1-3_ subunits are 94% identical between Gα_i1_ and Gα_i3_, 86% identical between Gα_i1_ and Gα_i2_, and 88% identical between Gα_i2_ and Gα_i3_ ^14^. These three members of the Gα_i_ subfamily have identical rates of single turnover GTP hydrolysis, but the GDP dissociation rate from Gα_i2_ is approximately two-fold faster than for the other two isoforms ^15^.

In terms of signaling specificity, all Gα_i_ subtypes inhibit various AC isoforms with similar potency and efficacy ^16^. For decades, AC was the only known effector of Gα_i_. Subsequently, a small number of proteins have been characterized as binding partners of Gα_i_: G protein-activated inwardly-rectifying potassium channels (GIRK) ^17–20 21^, epidermal growth factor receptor (EGFR), and growth factor receptor binding 2– associated binding protein 1 (Gab1) ^22^, although the biochemical and biological significance of these interactions is less well understood. Importantly, genetic deletion or inactivation of endogenous individual Gα_i_ isoforms have yielded evidence for differential function in primary tissues and organisms. For example, knockout of Gα_i2_ in mice results in exacerbated ischemic injury and cardiac infarction, while mice lacking Gα_i3_ saw an upregulation in Gα_i2_ and reduced injury ^21, 23–26^. Additionally, Gα_i2_ primarily promotes arrest and Gα_i3_ is required for transmigration and chemotaxis in mouse neutrophils ^27^, while Gα_i3_ activation downstream of CXCR3 has been shown to inhibit Gα_i2_ activation in murine activated T cells ^28^. These data strongly suggest that these isoforms serve non-redundant, unique functions, yet the biochemical basis for driving selective functionality has yet to be determined despite nearly three decades of research.

Recently, our laboratory identified PDZ-RhoGEF (PRG) as a novel, direct effector of Gα_i_ in an unbiased proximity interaction screen ^29^. Gα_i1_ binds and activates PRG in a nucleotide-dependent and receptor-dependent manner in cells. Gα_i3_ also activates PRG, but Gα_i2_ only weakly stimulates PRG. Here, we have interrogated the nature of the specificity of Gα_i_ subfamily members for PRG at the molecular level. In doing so, we have uncovered an atomic-level mechanism where the differences between Gα_i1_ and Gα_i2_ with respect to the ability to stabilize interactions between the HD and the Switch III region of the RLD results in weaker PRG engagement by Gα_i2_. Follow-up with unbiased proximity labeling coupled to tandem MS proteomics supports the idea that this mechanism extends beyond PRG interactions to multiple additional Gα_i_ targets. Overall, our studies support a model in which the strength and frequency of interactions between Gα_i_ Switch III and the HD control the ability to bind and activate PRG and other target proteins, differentiating Gα_i_ subfamily structure and function.

## Results

### Gα_i1_ more effectively activates and interacts with PRG than Gα_i2_

We have previously shown ^29^ that Gα_i1_ stimulates PRG and subsequent RhoA activation in a manner dependent on the activation state of Gα_i_. To mimic that GTP bound state of Gα_i_, a catalytic glutamine 204 was substituted with leucine which strongly inhibits GTP hydrolysis leading to constitutive GTP binding and activation ^7, 30–32^. Transient co-expression of Gα_i1_ Q204L (Gα_i1_ QL), PRG, and an SRE-luciferase plasmid that reports on RhoA activation in HEK293 cells (Fig. 1A) results in significant PRG activation (Fig. 1B). Gα_i2_ Q205L (Gα_i2_ QL) only weakly activates PRG activity in the same assay. Concentration-response experiments show a significant difference in the efficacy of PRG activation by Gα_i1_ QL and Gα_i2_ QL (Fig. 1C). This indicates that the difference is not due to differences in GTP binding since this would alter the potency of activation rather than efficacy. There is some variability in this assay with respect to the fold activation of PRG by Gα_i_ but the differences between Gα_i1_ and Gα_i2_ remain internally consistent within each assay set.

**Figure 1.**
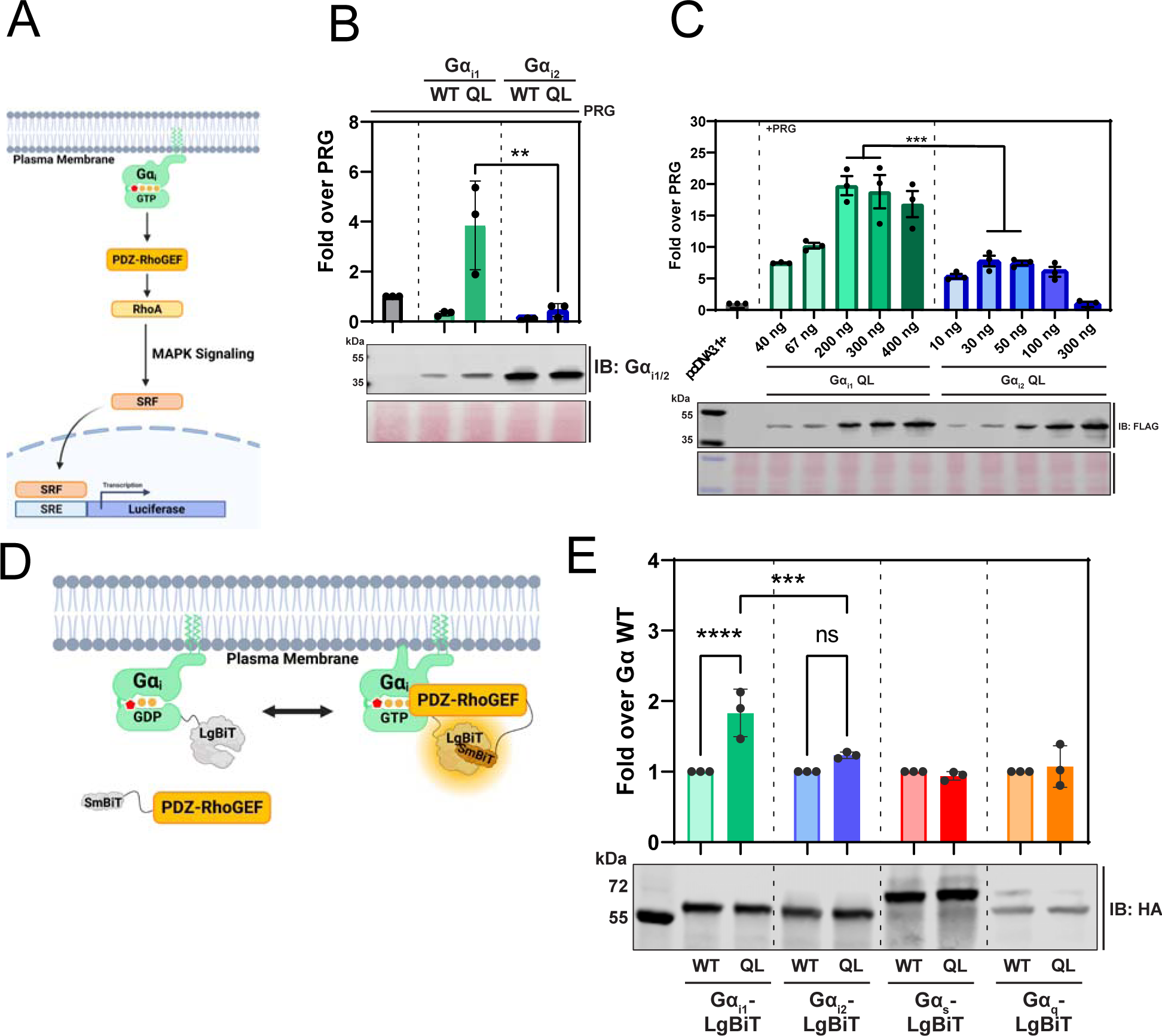
Gα_i1_ more efficiently interacts with PRG than Gα_i2_. **A)** Diagram of the SRE luciferase used to assess Gα regulation of PRG. HEK293 cells were co-transfected with control plasmid pcDNA 3.1 or Gα plasmids as indicated, PRG, and an SRE luciferase reporter plasmid. 24 h after transfection One-Glo luciferase reagent was added and luminescence was measured using a plate reader. **B)** Comparison of Gα_i1_ and Gα_i2_ which were transfected as indicated. All wells were transfected with PRG. Fold over PRG was calculated as the luminescent signal with Gα subunits co-transfected with PRG divided by the signal with PRG co transfected with control pcDNA 3.1 plasmid. **C)** Cells were transfected with the indicated amount of FLAG-Gα_i1_ QL or FLAG-Gα_i2_ QL adjusted to achieve equivalent expression as shown in the flag western blot shown in the bottom panel. To calculate the significance in the difference in maximal stimulation the values for 200 and 300 ng of Gα_i1_ plasmid were averaged and compared to the average of the 30 and 50 ng values for Gα_i2_. T-test *** P<0.001. **D)** Diagram of the Gα_i_-LgBiT complementation assay used with Gα_i_ fused to LgBiT and PRG with N-terminal fusion of SmBiT peptide natural peptide sequence (PRG-SmBiT). **E)** The indicated plasmids were co-transfected into HEK293 cells with PRG-SmBiT. 24 h after transfection cells were transferred into a 96 well plate and furimazine substrate was added for 15 min prior to measurement of luminescence in a plate reader. All experiments were performed with at least three biological replicates of assays performed in triplicate. Unless otherwise indicated data was analyzed with a one-way ANOVA with a Šídák post-test. ** P<0.01 and ****P<.0001.

To validate PRG-Gα_i_ interactions in cells, we performed a NanoBiT nanoluciferase complementation assay ^33^, in which the NanoLuc LgBiT was inserted after the αA helix in Gα subunits ^34^, and NanoLuc SmBiT was appended to the prior to the N-terminal Myc tag of myc-PRG (Fig. 1D). Coexpressing Gα_i1_ QL-LgBiT with SmBiT-PRG in HEK293 cells resulted in an increase in luminescent signal relative to Gα_i1_ WT-LgBiT, indicating a nucleotide-dependent interaction with PRG. This was not observed for QL variants in Gα_i2_, Gα_s_, or Gα_q_ (Fig. 1E). Together, these results show that Gα_i1_ interacts with, and activates PRG in a GTP-dependent manner, while Gα_i2_ is much less efficient in this interaction.

### Active Gα_i2_ QL BioID weakly engages the proximal interactome relative to Gα_i1_ QL BioID

Given their previously known functional overlap, the stark disparity between Gα_i1_ and Gα_i2_ in their ability to activate PRG prompted us to probe for further examples of selectivity between Gα_i_ subtypes. PRG was initially identified as a novel target of Gα_i1_ using unbiased BioID2 proximity labeling coupled to mass spectrometry. BioID2 functionalizes biotin releasing reactive biotinoyl-5’-AMP, which biotinylates proximal lysines within 20 nm ^35^. By comparing relative biotinylation by BioID2 fused to either Gα_i_ WT or Gα_i_ QL, we revealed the activated Gα_i_ proximity interactome. Here, we applied this approach to probe the relative interactomes of Gα_i1_ and Gα_i2_.

Briefly, HA-Gα_i1_ Q204L-BioID2 (Gα_i1_ QL-BioID), HA-Gα_i2_-BioID2 (Gα_i2_-BioID), and HA-Gα_i2_ Q205L-BioID2 (Gα_i2_ QL-BioID) were transiently transfected into HT1080 fibrosarcoma cells and incubated with biotin to allow labeling of proximal proteins by Gα_i_-BioID. After 24 hours of protein expression and biotin labeling, cells were lysed, biotinylated proteins were captured with streptavidin beads, and labeled with isobaric tandem mass tag (TMT) labels. Samples from all experimental groups were then analyzed via LC MS/MS (Fig. 2A). Proteins statistically significantly enriched in QL vs WT samples are considered proximal interactors. Volcano plots were generated for all the proteins identified with the statistical cutoffs for significance from two different comparisons, Gα_i1_ QL/Gα_i2_ WT (Fig. 2B top panel) and Gα_i2_ QL/Gα_i2_ WT (Fig. 2B bottom panel). We assumed that the Gα_i_ WT interactions would be similar between the two subtypes thus Gα_i2_ was used as a baseline for both plots. Validation of this assumption is discussed below.

**Figure 2.**
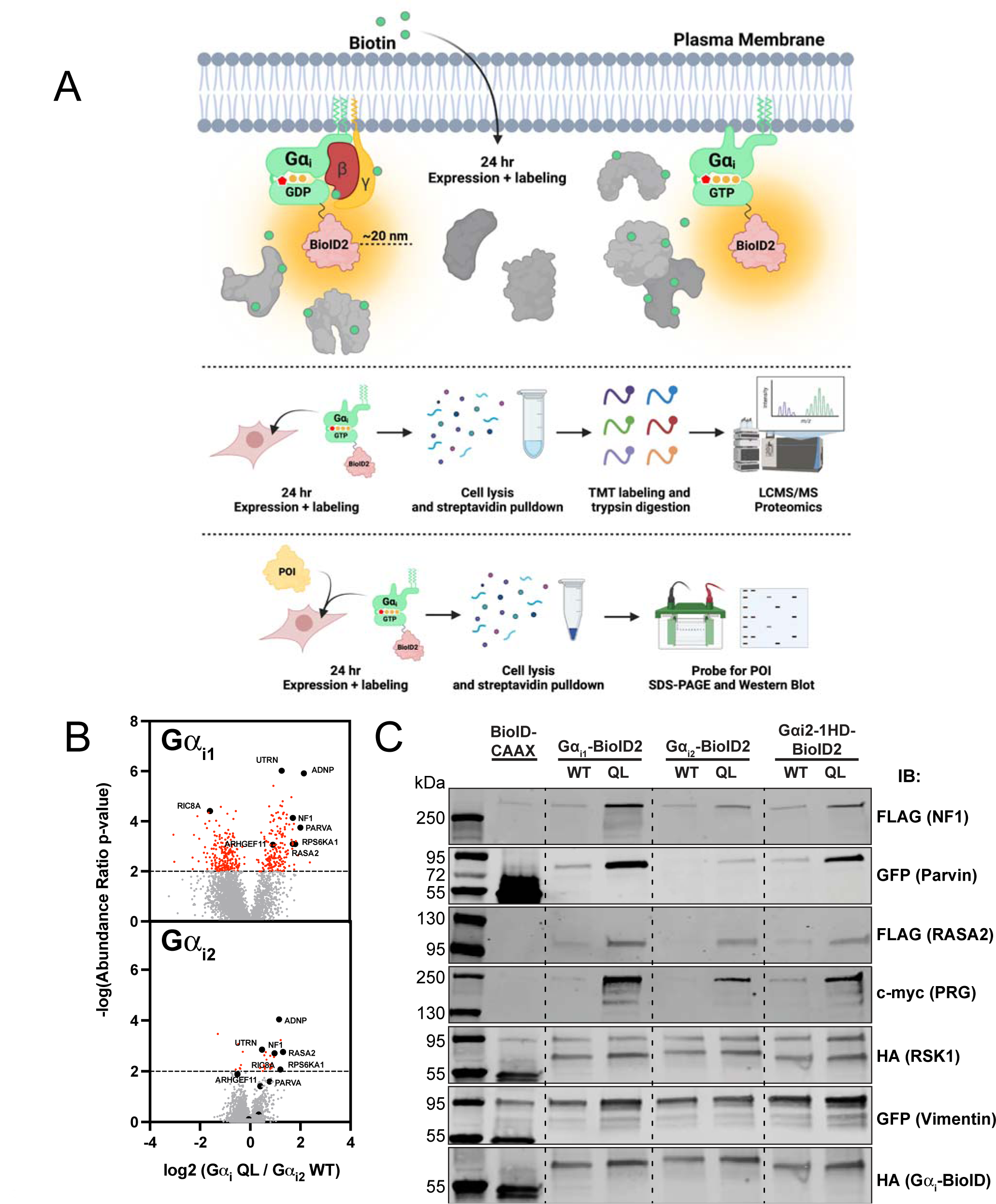
Active Gα_i2_ weakly engages the proximity interactome relative to Gα_i2_. **A)** Experimental outline for biotin proximity labeling assays. **B)** The indicated HA-Gα_i_-BioID2 constructs were transiently transfected into HT1080 cells, in triplicate for each condition for 24 h followed by isolation of biotinylated proteins and analysis by TMT Mass Spectrometry. To control for differences in overall biotinylation each sample was normalized based on the total spectral counts for all of the proteins identified (∼4000 proteins). Spectral counts were then analyzed as the ratio of samples transfected with the Gα_i_-QL plasmids relative to samples transfected with Gα_i2_ WT. The dashed line indicates a p value of 0.01 and all statistically significant proteins are colored in red. **C)** The indicated Gα_i_-BioID2 constructs were co-transfected with the indicated epitope-tagged protein into HEK293 cells. 24 h after transfection biotinylated proteins were isolated with streptavidin beads and the followed by western blotting to determine the amount of biotinylated target protein pulled down. Shown is a representative western blot of an experiment performed twice.

The identities and fold QL/WT enrichment levels for many hits for active Gα_i1_-BioID were consistent with those found in our previous screen ^29^. Notably, there are no significant observable differences in <u>identity</u> of most of the proteins enriched for interaction with active Gα_i1_ QL-BioID vs Gα_i2_ QL-BioID. However, the number of proteins identified that reached statistical significance [-log(abundance ratio p-value) ≥ 2.0] were markedly fewer in Gα_i2_ QL-BioID2 samples than in Gα_i1_ QL-BioID2 samples. This is largely because the Gα_i2_ QL-BioID2 / Gα_i2_ WT-BioID2 fold enrichment was generally lower than for Gα_i1_ QL BioID2. These data indicate a difference in overall signaling activity of Gα_i1_-GTP compared to Gα_i2_-GTP.

To confirm that these observations are not an artifact of the mass spectrometry analysis and that using Gα_i2_ WT as a baseline in both plots is valid, verification assays were performed with selected “hits” from the mass spectrometry that showed significant differences between Gα_i1_ QL and Gα_i2_ QL engagement. Epitope-tagged mammalian expression constructs were transiently co-expressed in HEK293 cells with either Gα_i1_-BioID, Gα_i1_ QL-BioID2, Gα_i2_-BioID2, Gα_i2_ QL-BioID2, or membrane-targeted BioID2 (BioID2-CAAX). Exogenous biotin was added for 24 hours, followed by a lysis and streptavidin bead purification. Captured biotinylated protein samples were run on SDS-PAGE and analyzed for pulldown via western blotting using antibodies against the respective affinity tags for the target proteins.

Proteins selected for analysis included several targets that were found in our previous report ^29^ and represent diverse signaling pathways: PDZ-RhoGEF, α-Parvin (Parvin), Vimentin, Ribosomal protein S6 Kinase A1 (RSK1), Neurofibromin 1 (NF1), and Ras p21 protein activator 2 (RASA2). Proteins including NF1, PRG, and Parvin showed selective enrichment in Gα_i1_ QL/WT over Gα_i2_ QL/WT (Fig. 2C, lanes 3-6). Vimentin and RASA2 showed only a slight preference for interaction with Gα_i1_ QL-BioID over Gα_i2_ QL-BioID, while RSK1 did not preferentially interact with either Gα_i1_ QL-BioID or Gα_i2_ QL-BioID over the WT-BioID variants. These results indicate that many of the proximal interactors found in the proteomic screen are reproducible in an orthogonal assay and are suitable for further analysis in their relationship to Gα_i_. Importantly, the results confirm that nucleotide-dependent interaction with these targets by Gα_i2_ is weaker than for Gα_i1_.

### Substitution of the Gα_i1_ helical domain (HD) into Gα_i2_ is sufficient to confer activation of PRG and enhances interactions with other targets

To understand the molecular determinants that drive specificity of activation of PRG by Gα_i1_, and perhaps by extension other targets, we mapped the amino acid differences between the Gα_i_ subfamily onto a crystal structure of Gα_i1_ bound to a GTP analogue, GPPNHP (PDB 1CIP). We previously reported that Gα_i3_ activates PRG, so we highlighted amino acids homologous between Gα_i1_ and Gα_i3_ but different from Gα_i2_ (33 residues) (Fig. 3A). The helical domain (HD) of Gα_i_ shows the region of greatest divergence between Gα_i_ subtypes (Figs. 3A and 4A), containing 21 of the differences between Gα_i1_/Gα_i3_ and Gα_i2_. As an initial approach, we substituted the entire HD of Gα_i1_ (residues 62-167) into the corresponding position in Gα_i2_, resulting in the chimeric Gα_i_ protein Gα_i2_-1HD (Fig. 3B). This chimera is expressed in HEK293 cells and functionally inhibits forskolin-dependent cAMP generation by adenylyl cyclase (Fig. S1A and B). Gα_i2_-1HD or Gα_i2_-1HD Q205L (QL) were then transfected into HEK293 cells in the SRE-luciferase reporter assay to examine their ability to activate PRG. Strikingly, Gα_i2_-1HD QL expression results in strong activation of PRG as compared to Gα_i2_ QL (Fig. 3C), indicating that the HD of Gα_i1_, when substituted into Gα_i2_, is sufficient to confer nucleotide-dependent activation of PRG.

**Figure 3.**
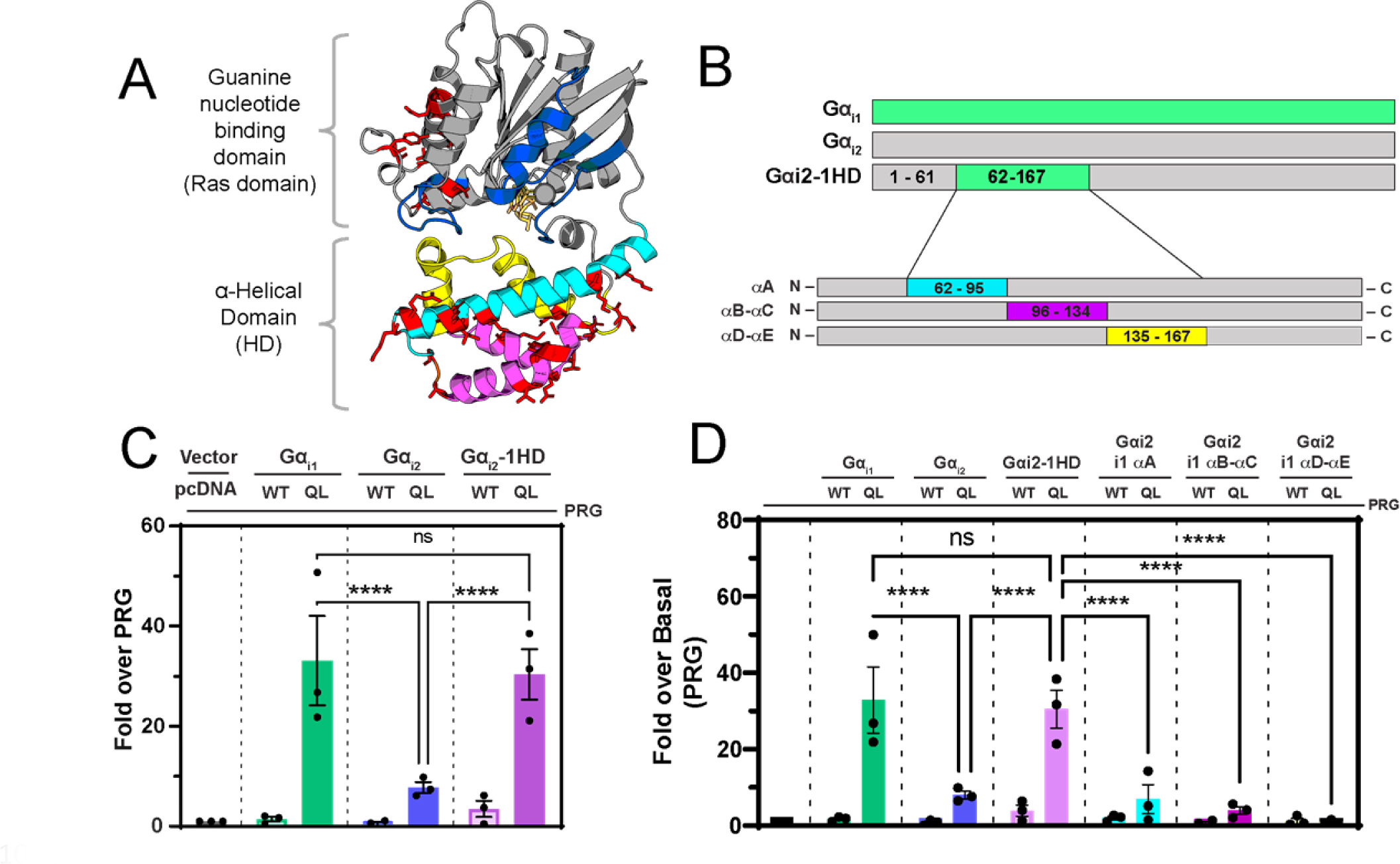
Substitution of the Gα_i1_ helical domain into Gα_i2_ partially restores activation of PRG. **A)** Diagrammatic representation of the Gα_i1_ structure. In cyan, magenta, and yellow are subdivisions of the helical domain. Switch I-III are in blue. Red stick amino acids are amino acids conserved between Gα_i1_ and Gα_i3_ but not Gα_i2_. PDB: 1CIP. **B)** Diagram of the constructs used in these experiments. **C) and D)** The indicated constructs were co-transfected with PRG and SRE-Luc and the assay was performed as in Fig. 1. Western blots for expression and cAMP assays are in Fig. S1 A,B,C and D. E) All SRE-luc experiments were performed with 3 biological replicates performed in triplicate. Data are +/-SEM analyzed by One-way ANOVA with Šídák post-test. **** P<0.0001.

**Figure 4.**
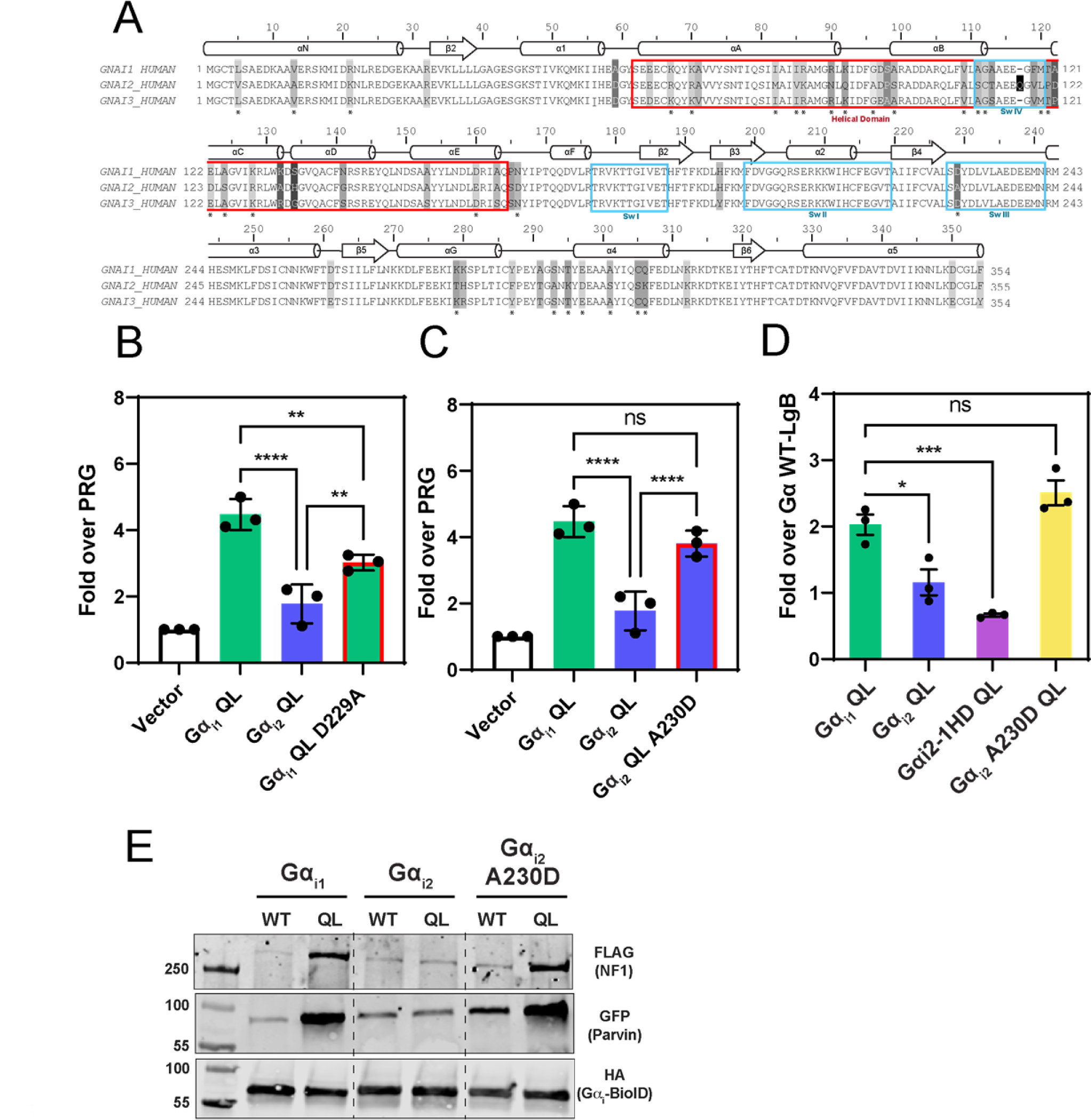
Gα_i1_ D229/Gα_i2_ A230^s4h3.3^ in the Ras-like domain is critical for differences in PRG activation. **A)** Alignment of human Gα_i1_, Gα_i2_, and Gα_i3_. Boxed in blue are the Gα_i_ switch regions. The helical domain is boxed in red. Starred (*) amino acids are identical in Gα_i1_ and Gα_i3_ but different in Gα_i2_. **B)** Mutation of Gα_i1_ D229^s4h3.3^ to the corresponding A in Gα_i2_ (A230^s4h3.3^) reduces the ability to activate PRG. **C)** Mutation of Gα_i2_ A230 to the corresponding D in Gα_i1_ (D229) enhances the ability of Gα_i2_ to activate PRG. **D)** Mutation of Gα_i2_ A230 to the corresponding D in Gα_i1_ (D229) enhances interactions between Gα_i2_-LgBiT and PRG-SmBiT in the luciferase complementation assay. **E)** Mutation of Gα_i2_ A230 to the corresponding D in Gα_i1_ (D229) enhances interactions with other proteins in the Gα_i_ proximity interactome. Shown is representative western blot for an experiment performed twice. All SRE-luc and complementation experiments were performed with 3 biological replicates performed in triplicate. Data are +/-SEM analyzed by One-way ANOVA with Šídák post-test; * P<0.05, ** P<0.01, *** P<0.001,**** P<0.0001.

To try to identify structural elements within the Gα_i1_ HD that confer PRG activation, the HD was subdivided into three segments consisting of 1) The Gα αA helix, 2) αB – αC helices, and 3) αD – αE helices. Each of these subdivisions of the Gα_i1_ HD were then substituted into their cognate positions in Gα_i2_ (Fig. 3B). Neither the αA helix nor the αB-αC helix subdivisions of Gα_i1_, when substituted into Gα_i2_, activate PRG in cells more than Gα_i2_ Q205L (Fig. 3D), but inhibited cAMP generation by adenylyl cyclase (Fig. S1C). The αD-αE substitution was deficient in the cAMP inhibition assay and could not be analyzed. These data suggest that Gα_i1_-mediated activation of PRG relies on some intrinsic property of the intact Gα_i1_ HD rather than one residue or a subset of residues within the Gα_i1_ HD. It is possible that the Gα_i1_ HD participates in direct binding interactions with PRG but may also confer specificity through interactions with of some component of the RLD in Gα_i_.

The striking increase in PRG activation observed with substitution of the Gα_i1_ HD into Gα_i2_ prompted us to test the interaction of these Gα_i2_ variants with other protein targets from the BioID proximity labeling screen. We tested multiple targets for activation-dependent labeling using the proximity labeling-dependent western blotting assay with the WT and QL versions of Gα_i1_, Gα_i2_ and Gα_i2_-1HD (Fig. 2C lanes 7,8, S1E). Substitution of the Gα_i1_ HD into Gα_i2_ partially rescues the QL-dependent labeling of some of these targets. Parvin shows the most striking rescue while NF1, PRG and vimentin show some degree of rescue. RASA2 which does not show a preference for Gα_i1_ vs. Gα_i2_ is not affected by the HD substitution. These data support the idea that the structural differences conferred by the HD of the Gα_i_ subunits are important for differences in general target engagement beyond PRG.

### Residue A230 in Gα_i2_ controls PRG activation and leads to enhanced proximity interactome engagement

Since we could not identify individual residues in HD that could confer PRG activation we hypothesized that the HD could be influencing contacts in other regions of Gα. In an existing co-crystal structure of Gα_13_ bound to the rgRGS domain of PRG ^36^, amino acids in the N-terminal portion of the PRG RGS domain bind at the Gα_13_ HD-RLD domain interface. We hypothesized that this paradigm may extend to PRG interactions with Gα_i1_ as well where the HD may cooperate with the Ras like domain to confer interactions with PRG. Based on this idea we individually substituted non-conserved residues (amino acids conserved between Gα_i1_ and Gα_i3_ but different in Gα_i2_, starred in Fig. 4A) from the Gα_i1_ RLD into Gα_i2_ and determined if they confer activation of PRG. The majority of the mutations either had no effect or reduced activation, however, substitution of Gα_i2_ A230^s4h3.3^ with Asp enables Gα_i2_(A230D) QL to activate PRG (Fig. 4B, Fig. S2A), while the reverse substitution of D229 to Ala in Gα_i1_ blunts PRG activation (Fig. 4C). The Gα_i2_ A230D substitution also confers the ability to interact with PRG in a nucleotide-dependent manner in the NanoBiT complementation assay in (Fig. 4D, Fig. S2B). We chose two of the other targets that show differential Gα_i1_ and Gα_i2_ engagement in the proximity labeling western blot assay, NF1 and Parvin, and performed the same assay comparing the QL versions of Gα_i1_-BioID2, Gα_i2_-BioID2 and Gα_i2_ A230D-BioID2 (Fig. 4E, Fig. S2C). The A230D substitution enhances the engagement of Gα_i2_ with these other targets. These data support the idea that the structural differences conferred by either the HD, or A230Gα_i2_/D229Gα_i1_ ^s4h3.3^, of the Gα_i_ subunits are important for differences in general target engagement beyond PRG. Additionally, the observation that these substitutions restore interactions previously identified in a Gα_i1_ BioID proximity labeling screen provides further evidence that these are in fact bona fide Gα_i_ interaction targets that remain to be further characterized physiologically.

### Gα_i1_ and Gα_i2_ sample distinct conformations

Examination of the static three-dimensional structure of Gα_i_**_1_** does not clearly indicate why substitution at the D229/A230^s4h3.3^ position, or substitution of the Gα_i1_ HD, would impact binding and/or activation of target proteins. This amino acid is near the GTP binding site but is not involved in interactions with the nucleotide, and the closest residue in the HD is 9Å away (Fig. 5A and B). To capture potential interactions that are not observable in the crystal structures, we performed molecular dynamics (MD) simulations with GTP-bound Gα_i1_ and Gα_i2._ We used the crystal structure of Gα_i_ (PDB ID:1CIP) as a starting structure for Gα_i1_ and generated a homology model of Gα_i2_ using this structure as a template. MD simulations were run for each system totaling to 5μs. Principal component analysis was used to characterize the dominant motions in Gα_i1_ and Gα_i2_. Principal Component 1 (PC1) in both proteins is rotation of the HD and RLD relative to one another (Movie S1 and 3). Principal Component 2 (PC2) is a domain “opening” motion where the HD opens relative to the RLD via the interdomain hinge region (Movie S2 and 4). We projected all the snapshots from MD simulations on these two principal components as shown in Fig. 5C. It is evident from Fig. 5C (top panel) that Gα_i1_ and Gα_i2_ sample distinct conformation clusters in these principal component coordinates. MD simulations show that even when bound to GTP, there is some degree of domain opening is possible in both Gα_i1_ and Gα_i2_ but the domain opening is more pronounced in Gα_i2_ compared to Gα_i1_. When these simulations were done for the mutants Gα_i1_ (D229A) the RLD-HD domain opening moved closer to that of Gα_i2_. Similarly, with the A230D substitution in Gα_i2_, moves closer to that of Gα_i1_ in the RLD-HD domain opening coordinate (Fig. 5C bottom panel).

**Figure 5.**
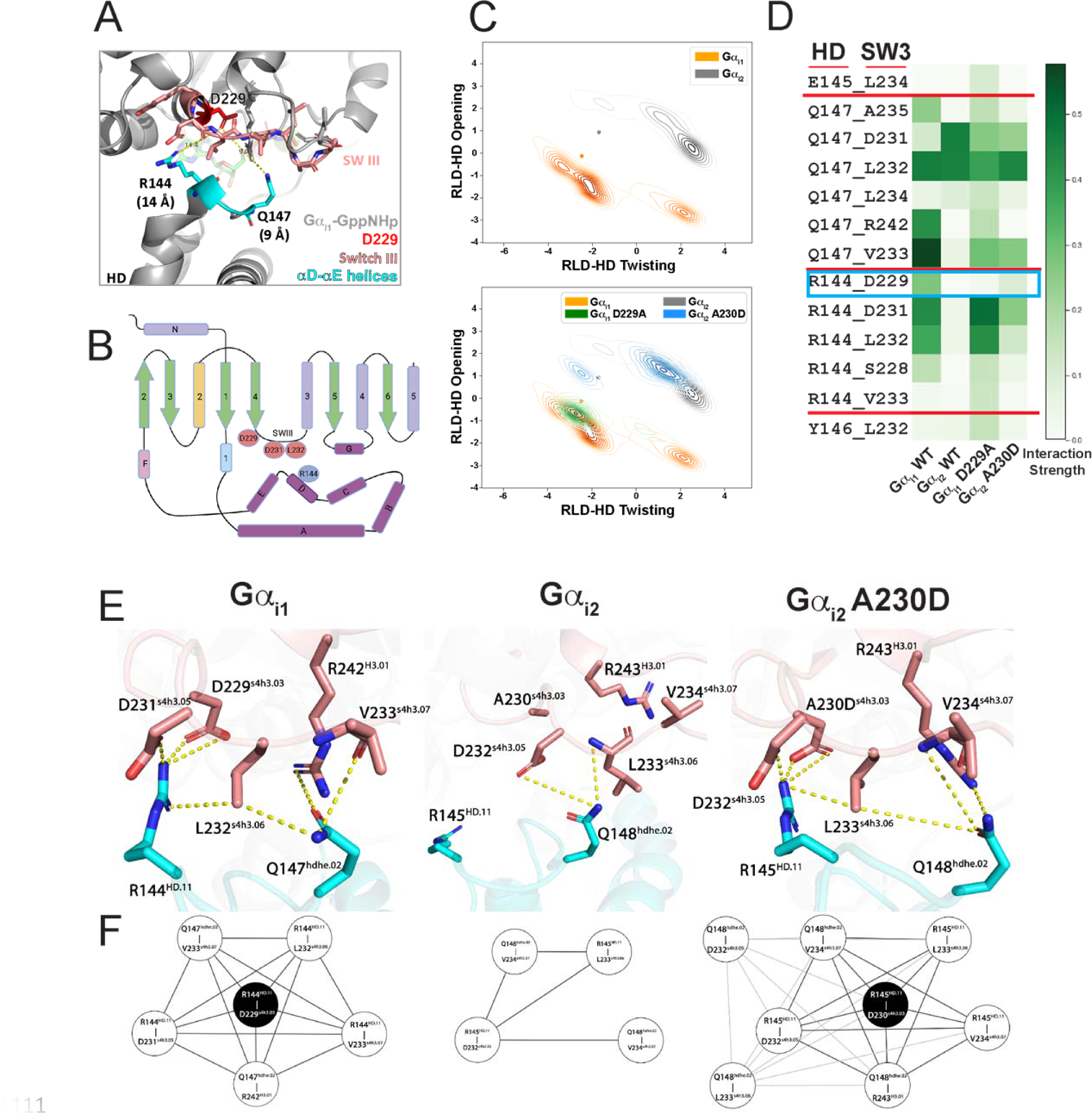
Molecular dynamics simulations and Bayesian network analysis reveal an interaction network that is not apparent in three dimensional crystal structures in the GTP bound state. **A)** Diagram of a structure of Gα_i1_-GTP showing the distance between D229 and the nearest HD residues. **B)** Ribbon representation of Gα subunit structure highlighting key amino acids at the Switch III-helical domain interface. **C)** Principal component analysis of Gα_i1_-GTP vs. Gα_i2_-GTP. **D)** Interaction frequency heat map of amino acid interactions between Switch III amino acids and amino acids in the HD comparing the GTP bound states of Gα_i1_, Gα_i2_, Gα_i1_ D229A, and Gα_i2_ A230D. **E)** Diagram of interdomain interactions involving D229 in Gα_i1_-GTP (top panel) and A230 in Gα_i2_-GTP (middle panel) and Gα_i2_-GTP A230D (right panel). **F)** Bayesian networks showing interdomain interactions driven by D229 and HD R144 in Gα_i1_-GTP(left panel), In Gα_i2_ A230 cannot interact with R145 weakening the overall interaction network (middle panel), Substitution of D for A230 in Gα_i2_-GTP leads to interactions with R145 stabilizing the interaction network between the HD and Switch III. Each node represents a contact made between the HD and Switch III, the thickness of the edge connecting the nodes indicates whether the edge was present in the Gα_i1_ network.

To understand the inter-residue interactions responsible for the differences in domain opening between these G protein subtypes, we analyzed the residues that make the interdomain contacts in the interface in all the MD snapshots. We observed differential interactions between residues in Switch III and the αD-αE region of the HD in Gα_i1_ compared to Gα_i2_ (Fig. 5D). In Gα_i1_, two key residues in the HD are involved in an interaction network at the HD-RLD interface, Q147^hdhe.2^ and R144^HD.11^. In our simulations during dynamic rotation of the HD-RLD interface, R144^HD.11^ dynamically interacts with residues D229^s4h3.3^, D231^s4h3.5^, L232^s4h3.6^, and S228^s4h3.2^ in the Switch III region of the RLD, interactions that are not evident in the crystal structure (Fig. 5E left). These interactions are largely absent in Gα_i2_ (Fig. 5E mid). In Gα_i2_, the cognate residue for Gα_i1_ D229 is A230, and substitution of A230 with D partially restores many of the interdomain residue interactions with Switch III that are absent in Gα_i2_ relative to Gα_i1_ (Fig. 5E right). Similarly, HD residue Q147^hdhe.2^ interacts more frequently with A235^s4h3.9^, R242^H3.1^, and V233^s4h3.7^ in Gα_i1_ than the cognate interactions in Gα_i2_. When Gα_i2_ A230^s4h3.3^ is substituted with D interactions between Q148^hdhe.2^ and V234^s4h3.7^ are strengthened, with other contacts are largely unaffected. This supports the idea that Gα_i1_ D229 stabilizes a network of interactions between the HD and RLD-Switch III that are lost in Gα_i2_ (Fig. 5D).

### Bayesian network models show that Gα_i2_ A230D mimics Gα_i1_ in RLD-HD interactions

As another approach, a fingerprint matrix of Switch III-HD residue contacts was constructed using data from the simulations. Bayesian Network Analysis was performed on this matrix, yielding a full Bayesian network (shown in Fig. S3 of Supporting Information) for these contacts in Gα_i1_ and Gα_i2_ and their mutants. Each node in this network model represents a residue interaction pair between RLD and HD. Nodes were then ranked by strength to understand their cooperativity ranking within the network. This analysis shows that interactions between D229^s4h3.3^ in the RLD and R144^HD.11^ in the HD forms the core of a cooperativity network involving multiple contacts in Switch III (Fig. 5F, left panel). This interaction network is disrupted in Gα_i2_ where the D229 cognate residue is alanine (Gα_i2_ A230) which cannot interact with the positively charged arginine (Gα_i2_ R145^HD.11^) (Fig. 5F, center panel). Substitution of A230 with D in Gα_i2_ restores a cooperative interaction network with Switch III (Fig.5F, right panel). This analysis supports the idea that in GTP-bound Gα_i1_, D229 at the base of Switch III forms an important contact with R144 in the HD that is not observed in crystal structures of Gα_i1_. This interaction supports a network of additional interactions between the HD and multiple amino acids in Switch III that constrain the conformation of Switch III. This network does not form in Gα_i2_, likely permitting Switch III to adopt conformations other than that seen in Gα_i1_, leading to lower-efficacy interactions with effectors that require Switch III for activation.

### PRG stimulation is dependent on interdomain stabilization of Gα_i_ Switch III

The simulation data indicate that an ionic interaction between D229 in the RLD and R144 in the HD centers an interaction network that controls the conformation of Switch III. Based on this we predicted that mutation of R144 to disrupt this interaction would reduce PRG activation by Gα_i1_. Gα_i1_ R144A reduces nucleotide-dependent PRG activation in cells, similar to that of Gα_i1_ D229A. When alanine is substituted for both D229 and R144, the same reduction is observed (Fig. 6A). Alanine substitution of cognate residue R145 in Gα_i2_ does not alter nucleotide-dependent PRG activation, but completely abolishes activation of PRG conferred by A230D (Fig. 6B). These experiments show that the D229-R144 interaction contributes to the ability of Gα_i1_ to activate PRG, and the ability to activate PRG conferred to Gα_i2_ by the A230D substitution is entirely dependent on the interdomain D230-R145 interaction.

**Figure 6.**
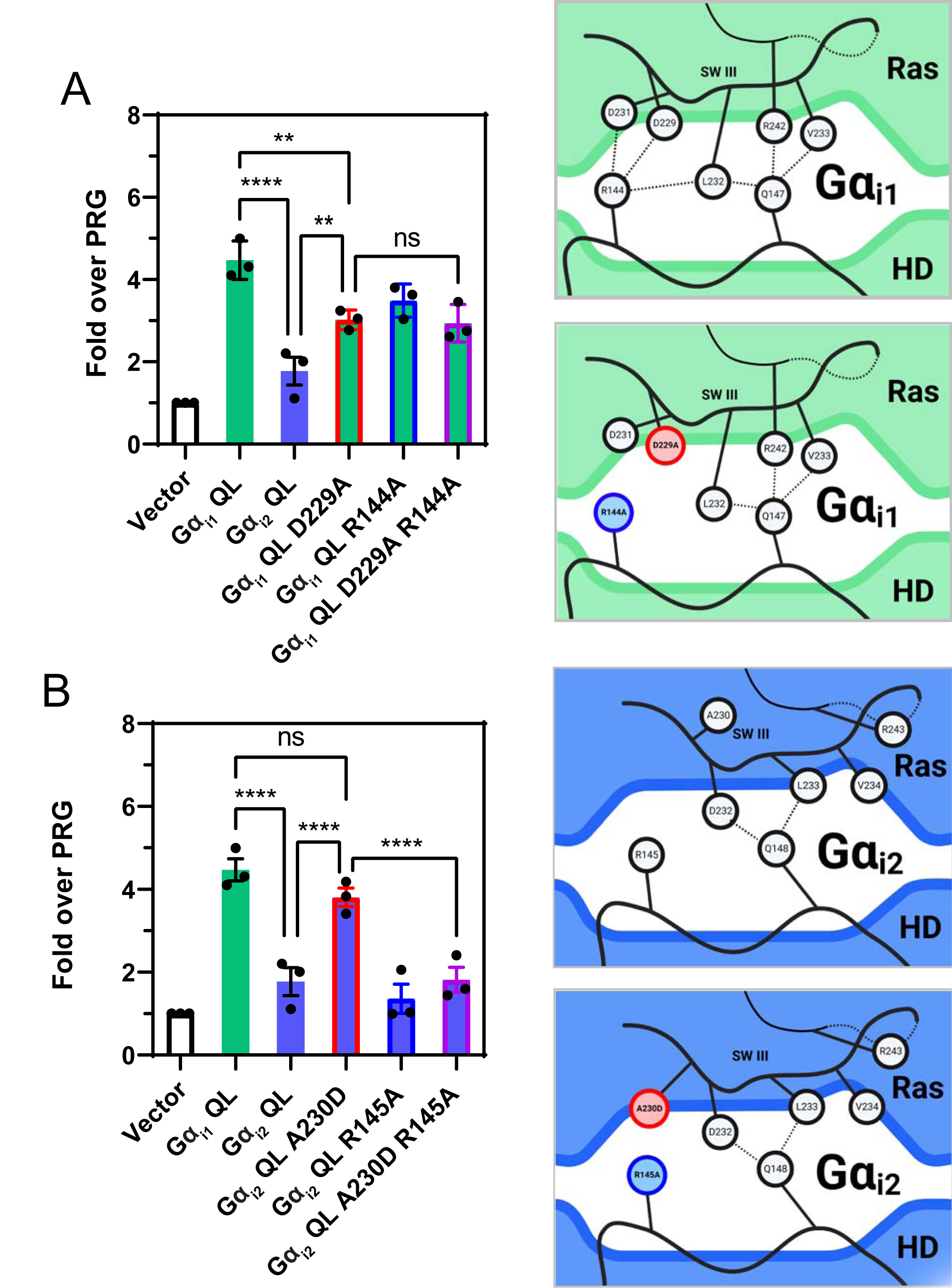
Gα_i1_ D229/Gα_i2_ A230 controls HD-RLD interdomain interactions. **A)** SRE luciferase assay showing PRG activation by QL versions of Gα_i1_, Gα_i2_, Gα_i1_ D229A, Gα_i1_ R144A, and Gα_i1_ D229A-R144A (left panel). The top right panel is a diagram of the WT Gα_i1_ interaction network. The bottom right panel is a diagram of the Gα_i1_ interaction network indicating the amino acid substitutions in red and blue. **B)** SRE luciferase assay showing PRG activation by QL versions of Gα_i1_, Gα_i2_, Gα_i2_ A230D, Gα_i2_ R145A, and Gα_i2_ A230D-R145A. The top right panel is a diagram of the WT Gα_i2_ interaction network. The bottom right panel is a diagram of the Gα_i2_ interaction network indicating the amino acid substitutions in red and blue. Experiments were performed with 3 biological replicates performed in triplicate. Data are +/-SEM analyzed by One-way ANOVA with Šídák post-test; * P<0.05, ** P<0.01, *** P<0.001,**** P<0.0001.

In the Ras-like domain are the switch regions including the Switch III loop. Switch III is critical for communication to the HD across the domain interface, and affects multiple aspects of Gα protein function, including effector recognition ^37, 38^ and receptor-mediated activation ^39^. In the cocrystal structure of Gα_13_ and PRG, Switch III makes multiple contacts with PRG. To test involvement of Switch III in Gα_i_-dependent PRG activation, we substituted Gα_i1_ Switch III residues D231 – A235 (DLVLA) to cognate Gα_s_ residues N254 – R258 (NMVIR) (Gα_i1_ SW3αS). Gα_i1_ SW3αS QL poorly activated PRG compared to Gα_i1_ QL in the SRE luciferase assay (Figs. 7A and B). To confirm that Gα_i1_ SW3αS retains activity Gα_i1_SW3αS was purified and compared with Gα_i1_ and Gα_i2_ for its ability to inhibit Gα_s_-stimulated adenylate cyclase. All three proteins were able to equally inhibit AC demonstrating that the Gα_i1_SW3αS chimera is functional (Fig. 7C). The loss-of-function mutations in Switch III along with the gain-of-function phenotype achieved by substitution of either Gα_i1_ RLD elements or HD elements provide evidence of cooperation between the RLD and HD stabilizing Switch III in a conformation needed for Gα_i_-mediated activation of PRG and other targets, but not inhibition of AC (Fig. 7D). This stabilization is lost in Gα_i2_.

**Figure 7.**
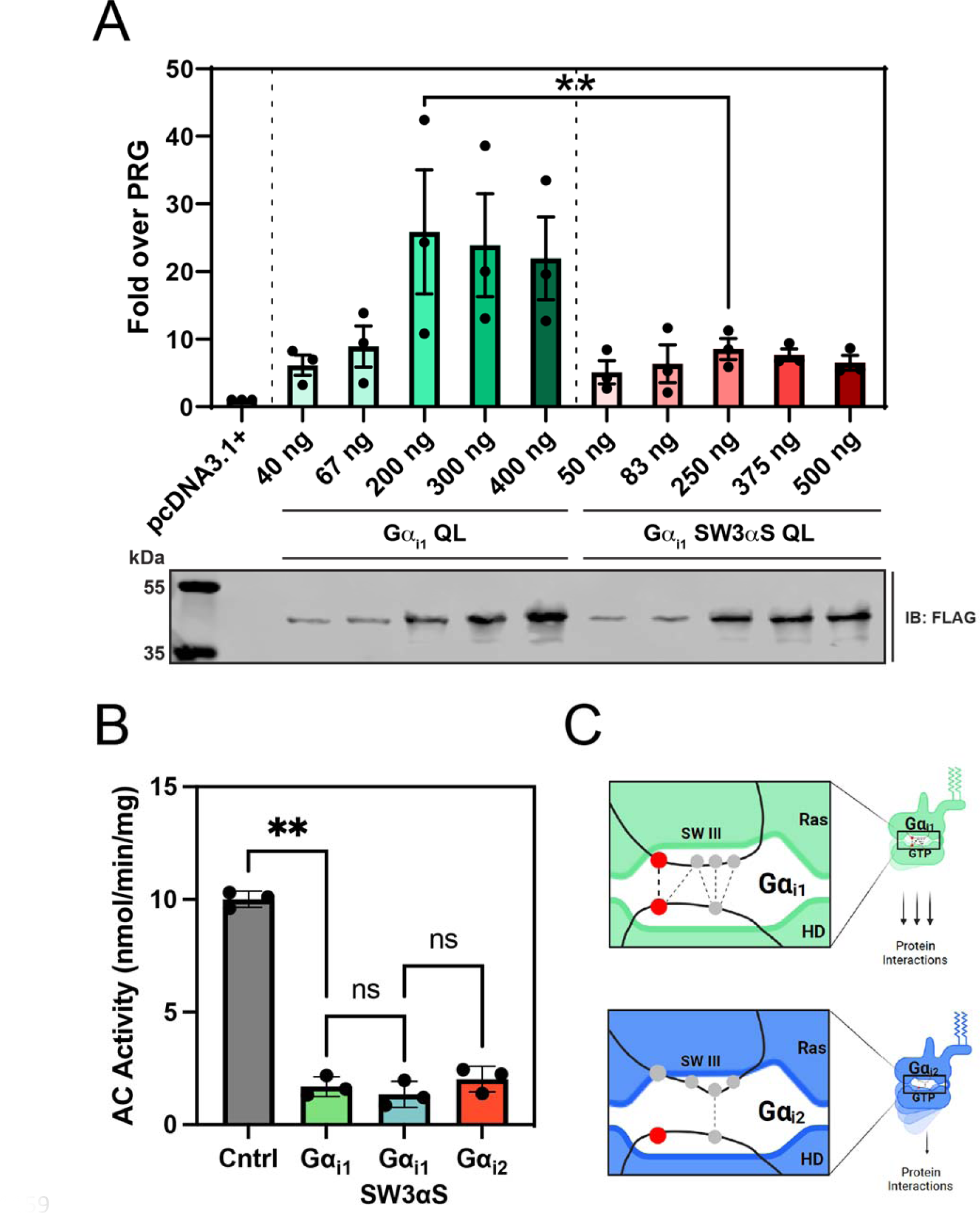
Gα_i1_ Switch III is critical for activation of PRG. **A)** Switch III amino acids in Gα_i1_ were substituted with the cognate amino acids in Gα_s_ and assayed for PRG activation using the SRE-luc assay. Experiments were performed with 3 biological replicates performed in duplicate. Data are +/-SEM analyzed by One-way ANOVA with Šídák post-test; * P<0.05, ** P<0.01, *** P<0.001,**** P<0.0001. **B)** Structural representation of active Gα_i1_ and active Gα_i1_ with Gα_s_ substitutions made in Gα_i1_ Switch III. Gα_i1_ is grey, the αD helix in the HD is shown in tan for orientation, Gα_i1_ Switch III residues are shown in green sticks, and the Gα_i1_ residues mutated to corresponding residues in Gα_s_ are in pink. PDB ID: 1CIP. **C)** Sf9 membranes expressing hAC6 were assayed in the presence of 10 mM MgCl_2_, 250 µM ATP and 30 nM Gα_s_·GTPγS in the absence and presence of 1µM myrGα_i1_, myrGα_i1_SW3αS, or myrGα_i2_. mean ± SD, n=3 performed in duplicate. **D)** Overall model of the interactions between the AHD and RLD domains of Gα_i_ subunits that modulate differential interactions between Gai subtypes and downstream proteins.

## Discussion

In this study, we provide evidence that Gα_i_-effector interactions are dependent on the strength and frequency of interaction between Switch III residues and the HD in the GTP bound state, and that these interactions differ between Gα_i_ subtypes. The data show that Gα_i2_ has fewer interdomain residue contacts, leading to weaker interactions between Switch III in the RLD and HD. Interruption of these contacts limits the ability of Gα_i_ to activate PRG. It is likely that stabilization of Switch III is central to this mechanism because Switch III conformational changes are dependent on the nucleotide binding state (GTP vs. GDP) while the conformation of the HD is generally not altered upon GTP binding. While we focused on PRG stimulation as a functional indicator of Gα_i_ specificity, the Gα_i_-BioID proximity labeling experiments demonstrate that there are global differences in GTP-dependent interactions between Gα_i_ subtypes and several novel targets, and that these differences depend on the same substitutions of residues from Gα_i1_ into Gα_i2_ that conferred specificity for PRG activation. This result indicates stabilization of interdomain interactions in the GTP state may play an unappreciated role in the downstream signaling function of Gα_i_ subunits, and a major role in differentiating Gα_i_ subtype function.

The involvement of the Gα_i1_ D229-R144 interaction and other additional interdomain contacts in stabilization of Switch III and effector interactions are supported by multiple key results. First, computational simulations show a dynamic interaction landscape where single substitutions affect the strength of other regional contacts. Second, substitution of either the Gα_i1_ HD or A230D into Gα_i2_ results in increased, GTP-dependent interaction with PRG and other protein targets compared to Gα_i2_ QL. Third, the effects of A230D in the RLD are completely abrogated if R145 in the HD is changed to alanine, strongly supporting the idea that this interdomain linkage is key to stabilizing the interface and Switch III such that it can interact with targets.

Position s4h3.3 (Gα_i1_ D229 and Gα_i2_ A230) is unique for the Gα_i_ subfamily in that the residue is different for each Gα family but is conserved within each family except Gα_i_. Amino acids at this position for each family include Ser in Gα_s_, Gly in Gα_o_ and Gα_z_, Ala in Gα_T_, and Glu in Gα_q/11_ and Gα_12/13_ (Fig. S5). A similar ionic lock mechanism for stabilization of Switch III through interdomain interactions is likely conserved in the Gα_q/11_ family and also Gα_13_, as Gα_i1_ R144^HD.11^ is conserved in these G proteins and could interact in a similar way with Glu at s4h3.3 in Switch III. Despite the similarities to other Gα subunits at these positions, the Gα_i_ subfamily seems unique in its intra-family effector specificity achieved by differentiation at s4h3.3 resulting in the presence or absence of the ionic lock.

RLD-HD interactions have classically been understood to be a regulator of nucleotide exchange ^12, 40–45^, with mutations at the interface intended to disrupt interactions leading to higher rates of GDP dissociation ^12^. Specifically, mutation of residue R144 in Gα_i1_ to an alanine is known to significantly increase the rate of GTPγS binding, presumably through the breaking of an interdomain interaction with L232 ^12^. In Gα_s_, substitution of residues in the Switch III loop to those of Gα_i2_ disrupt the ability of Gα_s_ to bind GTP in response receptor activation, but retains the ability to activate AC in response to GTPγS activation. Activation can then be restored by additionally substituting the Gα_s_ HD with Gα_i2_ residues ^39, 46^, demonstrating the importance of Gα isoform-specific interdomain communication for receptor dependent G protein activation.

Co-crystal structures of Gα subunits in each family have shown all non-RGS effectors binding to a common cleft between the α2 (Switch II) and α3 helices with no apparent direct involvement of Switch III ^36, 47–51^. On the other hand, mutagenic analysis Gα_q_-GRK2 interactions revealed involvement of both the HD and Switch III ^52^, an interaction not evident in the co-crystal structure of Gα_q_ with GRK2. As another example, Gα_T1_ binding to the autoinhibitory γ subunit of cGMP phosphodiesterase (PDEγ) is dependent on the presence of the HD ^53^, however the binding site of PDEγ is not in the HD but rather in the α2-α3 cleft ^51^. Crucially, mutation of a Switch III Glu to Leu abolishes PDE activation by Gα_T_, with no effects on nucleotide binding or hydrolysis^37^. A recent cryo-EM structure of the full cGMP PDE6 αβγ complex with transducin revealed the binding of PDEγ to the outer edge of the Switch III loop as well as the previously solved site in the α2-α3 cleft in Gα_T_-GTP ^54^. Thus, there is evidence for involvement of Switch III in effector engagement and our analysis reveals how two proteins with identical Switch III residues can have differences in target engagement efficacy.

While it remains untested how the lower efficacy of target engagement by Gα_i2_ relative to Gα_i1_ directly leads to distinct physiological roles, our findings are consistent with the notion that Gα_i2_ may in some situations act primarily to regulate AC and act as a scaffold and switch for Gβγ signaling, whereas Gα_i1_ or Gα_i3_ may perform these functions in addition to signaling to various Gα_i_-specific effectors. This is consistent with known roles for Gα_i2_ and Gα_i3_-mediated signaling events in neutrophils, where Gα_i2_ activation promotes cell arrest while and Gα_i3_ promotes migratory phenotypes ^27^. Eosinophils from Gα_i2_ whole-body knockout mice display enhanced chemotactic responses *in vitro* ^55^. The effects of activation of Gα_i2_ on neutrophil arrest in cells lacking Gα_i3_ are similar to those found by Gβγ activation alone ^56^. The physiological situation is likely to be more complex and this model cannot fully explain physiological specificity. For example, in murine atria, GIRK channel activity is differentially regulated by Gα_i2_ and Gα_i1_/Gα_i3_. Deletion of Gα_i2_ increases Gβγ-mediated basal and agonist-induced GIRK currents, while dual knockout of Gα_i1_ and Gα_i3_, which are known to bind and regulate GIRK, ablates basal and muscarinic agonist-induced GIRK activity ^57^. Nevertheless, it is probable that regulation of interdomain dynamics through the intramolecular interactions we defined play a significant role in physiological specificity.

In conclusion, we describe here a previously unknown mechanism of effector specificity between Gα_i_ subtypes. Switch III is stabilized by an interdomain interaction network with αD-αE residues in the helical domain, due in part to rearrangement of one non-conserved Gα_i_ Switch III aspartate that contacts a conserved arginine. This stabilization of Switch III not only confers specificity for activation of Gα_i1/3_ effector PDZ-RhoGEF, but for interaction with an array of additional protein targets, shedding light on a fundamental mystery of functional redundancy among this highly similar Gα protein family.

## Methods

### Plasmid cDNA constructs

BioID2 fused N-terminally with c-Myc tag and C-terminally with mVenus, followed by CaaX PM targeting motif (KKKKKKSKTKCVIM, derived from the C terminus of KRas), was a gift from S. Malik of the University of Rochester. C-terminally c-Myc–tagged full-length PRG cDNA construct in mammalian expression vector was a gift from J. Tesmer of Purdue University. The following plasmids were obtained from Addgene: mEmerald-parvin-C-14 (#54214), EGFP-vimentin-7 (#56439), HA-Gα_i_-BioID2 plasmids in pcDNA3.1+ were constructed as described previously ^29^.

All Gα clones in pcDNA3.1+ were obtained from the cDNA Resource Center. The sequences of the clones are available upon request.

All mutagenesis to Gα_i_ DNA constructs was accomplished using reagents, protocols, and guidelines from New England Biolabs Q5® Site-Directed Mutagenesis Kit (E0554S). Gαi2-1HD, all Gα**_i1_** HD subdivision constructs, and Gα_i_ N- and C-terminal substitutions were generated using reagents, protocols, and guidelines from New England Biolabs HiFi DNA Assembly Master Mix (E2621) and Cloning Kit (E5520).

In Gα_i1_, a FLAG epitope (DYKDDDDK) was inserted between Ala 121 and Glu 122 and flanked by a flexible linker (SGGGGS) on both sides of the insert. The FLAG epitope in Gα_i2_ was inserted in the same manner with the same linkers at the analogous position as Gα_i1_, between Asp 122 and Asp 123.

Gα_i1_ SW3αS-FLAG was generated using Q5 mutagenesis by substituting Gα_s_ residues N254 – R258 (NMVIR) into their cognate position in Gα_i1_, D231 – A235 (DLVLA) in FLAG-tagged Gα_i1_.

SmBiT-PRG was generated by inserting the SmBiT sequence (VTGYRLFEEIL) followed by a flexible linker (SGGGGS) onto the N-terminus of cMyc-PRG (cMyc: EQKLISEEDL), resulting in SmBiT-Linker-cMyc-PRG.

### Cell Culture

A293 and HT1080 cells were obtained from the American Type Culture Collection. A293 and HT1080 cells were grown supplemented in DMEM (Dulbecco’s modified Eagle medium) with 10% fetal bovine serum (FBS) (10437028, Gibco) and 100 U of penicillin/streptomycin (15140122, Gibco) at 37°C with 5% CO2. Trypsin-EDTA (25200056, Gibco) was used for cell passage.

### Reagents

The following primary and secondary antibodies were used: Gα_i1/2_ (anti-sera) ^58^, c-Myc (13-2500, Invitrogen), GFP (A11122, Invitrogen), HA (C29F4, Cell Signaling), FLAG (PA1-984B, Invitrogen). Streptavidin-IRDye800 was from LI-COR (925-32230). Primary antibodies were diluted in 3% bovine serum albumin (BSA) and 0.1% sodium azide and incubated with blots overnight at 4°C. Streptavidin-IRDye800 was incubated for 1 hour at room temperature. For secondary antibodies, goat anti-rabbit DyLight 800 (SA535571, Invitrogen) and goat anti-mouse IRDye 800CW (926-32210, LI-COR) were used at 1:10,000.

### NanoBiT Luciferase Complementation Assay

6.0 x 10^5^ HEK293A cells were seeded in poly-D-lysine coated 6-well plates (Fisher FB012927). Immediately after plating, HA-Gα-LgBiT constructs and SmB-cmyc-PDZ-RhoGEF were co-transfected using a 1:3 mass to volume ratio of DNA to Lipofectamine 2000 (Invitrogen). After 24 hours, transfection media was aspirated and cells were gently washed once with 1 mL warm PBS. The PBS was discarded, 200 μL trypsin solution was added, and the plate was incubated at 37°C and 5% CO_2_ for 5 mins. Following incubation, 800 μL of warm 1X HBSS was added to each well, and the detached cells were aspirated and dispensed into new 15 mL conical tubes. Cells were then pelleted by centrifugation at 250 x g for 5 mins at RT. After carefully aspirating the supernatant, each pellet was resuspended in 1 mL warm HBSS, and cell number in each suspension counted. Cell suspensions were centrifuged once more at 250 x g for 5 mins at RT and resuspended in warm 10 μM furimazine in HBSS, 1% DMSO. 5 x 10^4^ cells were distributed to each well in a 96-well plate; samples were analyzed with six technical replicates. The sample plate was incubated at 37°C for 15 mins, followed by a luminescence measurement in each well.

### SRE-Luciferase Reporter Assay

#### 96-well format

4.5 x 10^4^ HEK293A cells were seeded in poly-D-lysine coated 96-well plates (Greiner 655983). Cells were transfected with the following plasmids and amounts per well: 25 ng SRE-Luc reporter (E134A, Promega), 75 ng Gα_i_ or Gα_i_ QL in pcDNA3.1+, 2.5 ng cmyc-PRG unless otherwise indicated. Minor adjustments in added DNA were made to equalize expression of Gα_i_ subunits based on western blotting of Flag tagged constructs. In these cases, empty pcDNA3.1+ vector supplemented to equalize total DNA added per well. Transfection took place immediately after seeding with a 1:3 mass to volume ratio of DNA to Lipofectamine 2000 (Invitrogen). Twelve hours after transfection, the media was replaced with 75 µL of serum-free media. Twenty-four hours after transfection, 75 µL (1:1 volume) of One-Glo reagent (E6110, Promega) was added to each well and incubated for 10 min at room temperature. The luminescence signal was measured using Varioskan LUX multimode microplate reader (Thermo Fisher Scientific).

#### 24-well format

The SRE-Luc reporter assay was also performed nearly identically in 24-well plates, which offered better well-to-well consistency for technical replicates. 1 x 10^5^ HEK293A cells were seeded in poly-D-lysine coated 24-well plates. One hundred ng SRE-Luc reporter (E134A, Promega), 300 ng Gα_i_ or Gα_i_ QL in pcDNA3.1+, and 5 ng cmyc-PRG DNA were transfected into each well except in Gα_i_ titration experiments, where reduced Gα_i_ DNA was substituted with empty pcDNA3.1+. Transfection took place immediately after seeding with a 1:3 mass to volume ratio of DNA to Lipofectamine 2000 (Invitrogen). Twelve hours after transfection, the media was replaced with 250 µL of serum-free media. Twenty-four hours after transfection, 250 µL (1:1 volume) of One-Glo reagent (E6110, Promega) was added to each well and incubated for 10 min at room temperature. The luminescence signal was measured using Varioskan LUX multimode microplate reader (Thermo Fisher Scientific). We found that the fold differences in activation by Gα_i_ were lower in the 24 well format but that the technical replicates were more reliable.

### GloSensor cAMP Assay

4.5 x 10^4^ HEK293A cells were seeded in poly-D-lysine coated 96-well plates (Greiner 655983). Cells were transfected with the following plasmids and amounts per well: 50 ng GloSensor −20F cAMP plasmid (E1171, Promega), 125 ng Gα_i_ or Gα_i_ QL in pcDNA3.1+. In Gα_i_ titration experiments, DNA was supplemented with empty pcDNA3.1+ vector. Transfection took place immediately after seeding with a 1:3 mass to volume ratio of DNA to Lipofectamine 2000 (Invitrogen). Twenty-four hours post-transfection, the media was discarded and the cells were loaded with 75 µL 0.5 mg/mL D-Luciferin (L2916, Sigma Aldrich) in Leibowitz’s L-15 medium followed by incubating for 2 hours at 37°C and 5% CO_2_. Plates were removed from the incubator and equilibrated at room temperature. Forskolin was then added to give a 1 mM final concentration and luminescence was measured at 15 min in a plate reader.

### Western blotting

Samples in 1X Laemmli sample buffer were resolved on 4-20% gradient Mini-PROTEAN TGX gels (4561094, Bio-Rad), transferred to nitrocellulose membranes (Pall 66485), and stained with Ponceau S (141194, Sigma Aldrich). Membranes were blocked with 3% bovine serum albumin (141194, Sigma Aldrich) in TBST (0.1% Tween-20 in 20 mM Tris pH 7.5 + 150 mM NaCl) at room temperature (RT) for 30 min with constant agitation. Primary antibodies were applied for 2 hours at RT or overnight at 4°C. After three RT washes with TBST at 5 min each, secondary antibodies were applied for 1 hour. Membranes were imaged on an Odyssey Infrared Imaging System (LI-COR Biosciences).

### BioID2 proximity labeling and tandem mass spectrometry analysis

HT1080 cells at passage number up to 15 were used for proximity labeling experiments. Cells were plated into 175 cm^2^ flasks at a density of 5.5 × 10^6^ cells per flask. The next day, media was replaced with 35 mL of DMEM containing 50 µM biotin and 10% FBS. Each flask was transfected with 8 µg of plasmid encoding BioID2-fused Gα_i_ construct and 4 µg of YFP cDNA. A total of 0.6 µL of Viromer Red (VR-01LB-00, Lipocalyx, Germany) reagent was used per 2 µg of cDNA for transfection, resulting in ∼80 to 85% transfection efficiency. Twenty-four hours after labeling and transfection, the labeling medium was decanted, cells were washed twice with 1× PBS, and harvested at 4000 x g for 10 min. This step was repeated twice using 1× PBS to recover the maximum number of cells. The supernatant was aspirated, and pellets were flash-frozen and stored at −80°C until further use.

All stock solutions used for streptavidin pulldown were freshly prepared, except lysis buffer. Low protein binding tubes (022431081, Eppendorf) were used for sample preparation. Frozen pellets were lysed in 1 mL of ice-cold lysis solution (composition described above) for 10 min on ice and incubated with 125 U of benzonase with end over-end rotation at 4°C for 20 min. A total of 0.3% SDS was added to lysates, which were incubated for another 10 min at 4°C. Lysates were centrifuged at 15,000 x g for 15 min. The supernatant was transferred to fresh tubes, and the total protein concentration was measured using Pierce 660 nm protein assay reagent. A total of 5% of lysates, adjusted for protein concentration, was reserved to analyze the biotinylation in inputs. The remaining lysates were incubated with 500 µL of Pierce streptavidin magnetic beads slurry per sample in an end-over-end rotator at 4°C overnight. Beads were washed twice with modRIPA buffer [modRIPA: 50 mM tris, 150 mM NaCl, 0.1% SDS, 0.5% sodium deoxycholate, and 1% Triton X-100 (final pH 7.5)] and once with four different solutions: 1 M KCl, 0.1 M Na_2_CO_3_, 2% SDS [in 50 mM tris (pH 7.5)], and 2 M urea [in 10 mM tris (pH 8.0)]. Beads were washed twice with 1× PBS and were flash-frozen and stored at −80°C until further processed for MS.

### BioID2 proximity labeling and immunoblot analysis

1.5 x 10^6^ HEK293A cells were seeded in a poly-D-lysine coated 10 cm plate. The next day, media was replaced with 10 mL DMEM +10% FBS and biotin was added to 50 µM. Cells were transfected with 3 µg of either BioID-CAAX or one of the Gα_i_-BioID2-HA constructs in pcDNA3.1+, in addition to 3 µg of one of the effectors of interest (cmyc-PRG, V5-ADNP, RASA2-FLAG, mEmerald-Parvin, RSK1-HA, or GFP-Vimentin). DNA complexes were added to Lipofectamine 2000 solutions with a 1:3 mass:volume ratio (18 µL per plate). After 24 hours of expression and labeling, the medium was decanted, cells were rinsed twice with 5 mL of ice cold 1X PBS, scraped off of the plate, and pelleted at 4°C and 4000 x g for 10 min. The supernatant was aspirated and the cell pellets were flash-frozen with liquid N_2_ and stored at −80°C until processed via IP.

For the IP, 500 µL ice cold modRIPA was used to resuspend cell pellets. Lysis using benzonase and SDS proceeded as above. Lysates were centrifuged for 15,000 x g for 15 min at 4°C, and protein concentration was measured using Pierce 660 nm protein assay reagent. After equalizing for protein concentration, 20 µL of each sample volume was retained as an input sample. Five hundred µL of each equalized sample was added to 170 µL of Pierce streptavidin magnetic bead slurry and rotated end-over-end at 4°C for at least 2 hours to capture biotinylated proteins. Beads were washed three times with ice cold modRIPA and once more with cold 1X PBS. Beads were then resuspended in 100 µL 1X PBS, and 4X Laemmli sample buffer was added to 1X final concentration. Beads were boiled for 10 min at 95°C, and the supernatant was analyzed by western blot using anti-HA (1:2000) for Gα_i_-BioID2-HA and the corresponding antibody for each protein of interest [cmyc-PRG – anti-cmyc (1:2000), V5-ADNP – anti-V5 (1:1000), RASA2-FLAG – anti-FLAG (1:1000), mEmerald-Parvin – anti-GFP (1:1000), RSK1-HA – anti-HA (1:2000), or GFP-Vimentin – anti-GFP (1:1000)].

### Protein digestion and TMT labeling

On-bead digestion followed by liquid chromatography–tandem MS (LC-MS/MS) analysis was performed at the MS-based Proteomics Resource Facility of the Department of Pathology at the University of Michigan. Samples were reduced (10 mM dithiothreitol in 0.1 M triethylammonium bicarbonate (TEAB) at 45°C for 30 min), alkylated (55 mM 2-chloroacetamide at room temperature for 30 min in the dark), and subsequently digested using a 1:25 ratio of trypsin (V5113, Promega):protein at 37°C with constant mixing. A total of 0.2% trifluoroacetic acid was added to stop the proteolysis, and peptides were desalted using a Sep-Pak C18 cartridge (WAT036945, Waters Corp). The desalted peptides were dried in a vacufuge and reconstituted in 100 μl of 0.1 M TEAB. A TMT10plex Isobaric Label Reagent Set plus TMT11-131C Label Reagent kit (A37725, Thermo Fisher Scientific) was used to label each sample per the manufacturer’s protocol. The samples were labeled with TMT 11-plex reagents at room temperature for 1 hour. The reaction was quenched by adding 8 µL of 5% hydroxylamine for 15 min and dried. An offline fractionation of the combined sample into eight fractions was performed using a high pH reverse-phase peptide fractionation kit, as per the manufacturer’s protocol (84868, Pierce). Fractions were dried and reconstituted in 12 µL of 0.1% formic acid/2% acetonitrile for LC-MS/MS analysis.

### LC-MS analysis

An Orbitrap Fusion (Thermo Fisher Scientific) and RSLC Ultimate 3000 nano-UPLC (Dionex) were used to acquire the data. For superior quantitation accuracy, we used multinotch-MS3 ^59^. Two microliters of each fraction was resolved on a nanocapillary reverse-phase column (75 µm internal diameter by 50 cm; PepMap RSLC C18 column, Thermo Fisher Scientific) at a flowrate of 300 nL/min using 0.1% formic acid/acetonitrile gradient system (2 to 22% acetonitrile in 110 min; 22 to 40% acetonitrile in 25 min; 6-min wash at 90% acetonitrile; 25 min re-equilibration) and directly sprayed onto the Orbitrap Fusion using EasySpray source (Thermo Fisher Scientific). The mass spectrometer was set to collect one MS1 scan [Orbitrap; 120,000 resolution; AGC target 2 × 105; max IT (maximum ionization time) 50 ms] and data-dependent, “Top Speed” (3 s) MS2 scans [collision-induced dissociation; ion trap; NCE (normalized collision energy) 35; AGC (automatic gain control) 5 × 103; max IT 100 ms]. For multinotch-MS3, the top 10 precursors from each MS2 were fragmented by high energy collisional dissociation (HCD), followed by Orbitrap analysis (NCE 55; 60,000 resolution; AGC 5 × 104; max IT 120 ms, 100 to 500 mass/charge ratio scan range).

### Purification of Gα_i_ subunits

C-terminally hexahistidine tagged Gα_i_ subunits and chimeras were co-expressed with *N*-myristoyltransferase in E. coli as previously described ^60^. Proteins were purified using Ni-NTA chromatography using a gradient from 0-200 mM imidazole which resulted in proteins of greater than 90% purity. Myristoylation was confirmed by analyzing molecular weights on SDS-PAGE and G protein nucleotide binding activity was assessed using [^35^S]-GTPγS binding. All proteins had 20-40% nucleotide binding activity.

### Adenylyl Cyclase Activity Assays

Membranes from Sf9 cells expressing hAC6 (10 μg per reaction) were assayed for adenylyl cyclase activity as described ^61^. Purified and GTPγS-activated myristoylated Gα_i1_SW3αs, Gα_i1_ and Gα_i2_ were preincubated with membranes for 5 min on ice. Gα_s_·GTPγS (30 nM final) was added and preincubated for 5 min on ice prior to the start of the assay (10 min at 30 C). Reactions were stopped with 0.2N HCL and cAMP was detected by enzyme immunoassay (Assay Designs).

### Generating structural models and molecular dynamics simulations

The structural model of monomeric GTP-bound Gα_i1_ and Gα_i2_ protein with Mg^2+^ ion was built using the monomeric GTP bound rat Gα_I1_ crystal structure (PDB ID: 1CIP) as template and using the homology modeling method in the Prime module of Maestro software from Schrodinger (https://www.schrodinger.com/products/maestro). The GNP present in the original crystal structure was converted to GTP using Maestro edit panel. Point mutations to generate the structures of Gα_i1_^D229A^ _and Gαi2_^A230D^ were performed using Maestro Biologics suite. The side chain packing was done for all the residues within 5Å of the mutated residue position including the mutated residues using Maestro Prime suite. All structures further underwent energy minimization using conjugate gradient method with a convergence cutoff of 0.1kcal/mol/Å. Input files for molecular dynamics simulations were generated using CHARMM-GUI ^62^. Each monomeric Gα_i_ protein was solvated in explicit TIP3P water molecules in a cubic box (9.0nm x 9.0nm x 9.0nm) with 0.15M of potassium chloride to mimic the physiological condition. We used GROMACS software ^63^ (Version 2021.3) with all-atom CHARMM36 force field ^64^ to perform molecular dynamics simulations. MD simulations were performed at 310°K coupled to a temperature bath with a relaxation time of 0.1ps ^65^. Pressure of the systems was calculated with molecular virial and was held constant by a weak coupling to a pressure bath with a relaxation time of 0.5ps. Equilibrium bond length and geometry of water molecules were constrained using the SHAKE algorithm ^66^. The short-range electrostatic and van der Waals interactions were estimated every 2fs using a charged group pair list with cutoff of 8Å between centers of geometry of charged groups. Long-range van der Waals interactions were calculated using a cutoff of 14Å and long-range electrostatic interactions were treated with the particle mesh Ewald method ^67^. Temperature was kept constant at 310°K by applying the Nose-Hoover thermostat ^68^. Desired pressure for all systems were achieved by using Parrinello-Rahman barostat with a pressure relaxation time of 2ps ^69^. Before production runs, all system were subjected to a 5000-step steepest descent energy minimization to remove bad contacts^70^. After minimization, the systems were heated up to 310°K under constant temperature-volume ensemble (NVT). The simulations were saved every 200ps for analysis. The protein, Mg^2+^ ion, and nucleotide were subjected to positional constraints under a harmonic force constant of 1000 kJ/(mol*nm^2^) during the NVT step while solvent molecules were free to move. The systems then were further equilibrated using a constant pressure ensemble (NPT), in which the force constant is applied to the protein, Mg^2+^ ion, and nucleotide were gradually reduced from 5kJ/(mol*nm^2^) to zero in six steps of 5ns each. An additional 50ns of unconstraint simulation was performed, making it a total of 80ns NPT equilibration prior to production runs. We performed five production runs of 1000ns each using five different initial velocities for every system. Therefore, we had 5μs long MD trajectories for both WT and mutant systems of Gα_i1_ and Gα_i2_ protein.

### Principal Component Analysis and representative structures

The last 600ns of five independent molecular dynamics simulation runs were merged into one concatenated trajectory for each system. Two merged trajectories were further created based on the concatenated trajectories: one contains the WT Gα_i1_ and Gα_i2_ trajectories, and the other contains all four trajectories. Principal component analysis was performed on each merged trajectory using the gmx covar module of GROMACS with covariance matrix of C alpha atoms of all residues. The first two principal components (PC1 and PC2) of every system were extracted using gmx anaeig module of GROMACS and imported into Python as a data-frame using the Pandas package. Kernel density estimation maps were generated using Python Seaborn package (version 0.9.0) and plotted using Python Matplotlib package.

### Representative structure extraction

Using Get-contact data (see previous), frame numbers in Gα_i2_^A230D^ trajectory that have contacts between R145 and D230 were recorded. The corresponding frames were extracted from the trajectory using gmx trjconv module of GROMACS. The representative structure of Gα_i1_ was used as template, and the root-mean-square deviation (RMSD) values of the extracted Gα_i2_^A230D^ frames were calculated using gmx rms module: C alpha atoms were selected for both alignment and calculation. The frame with the smallest RMSD value was selected as the representative structure for Gα_i2_^A230D^ system.

### Calculating the fingerprints of pairwise interactions between AHD and switch III domain of G protein

The analysis of the landscape of pairwise intermolecular residue contacts between AHD domain and switch III region of Gα_i_ with MD simulations using the “getcontacts” python script library (https://www.github.com/getcontacts). This was utilized to identify various types of contacts, including salt-bridges (<4.0 Å cutoff between anion and cation atoms), hydrogen bonds (<3.5 Å cutoff between hydrogen donor and acceptor atoms, <70° angle between donor and acceptor), van der Waals (<2 Å difference between two atoms), pi-stack contacts (<7.0 Å distance between aromatic centers of aromatic residues, <30° angle between normal vectors emanating from aromatic plane of each residue), and cation-pi contacts (<6.0 Å distance between cation atom and centroid of aromatic rink, <60° angle between normal vector from aromatic plane to cation atom). To conduct the analysis, the MD simulation trajectories were concatenated into 1μs ensembles and stored as xtc coordinate files. Subsequently, water and ions were stripped from the trajectory files utilized for the contact analysis, and atom selection groups were matched with the relevant amino acid residues for each protein domain. In-house python scripts were used to perform one-hot encoding to generate a binary fingerprint for each simulation. The one-hot encoding represented the presence of a contact between two residues in a particular frame with “1” and its absence with “0”.

### Bayesian Network Analysis

Binary fingerprints of residue contact pairs were analyzed to understand their interdependent interactions using BNOmics, software developed for Bayesian network analysis. Separate BNs were first constructed for each G protein type. Heuristic network model selection search ^71^ was carried out with 50 random restarts, to ensure convergence. Bayesian networks of contact fingerprints have residue pairs as nodes and the edge weight between the nodes correlates with the dependency between them. As a measure of contact pairs’ connectivity, the network property of node strength was used - the total sum of edge weights belonging to this node. After sorting the residue pairs from highest node strength to the lowest, the top 25 percentile of them was compared between different G protein types. Graphical representation of these nodes and their interconnections were demonstrated using network visualization software Cytoscape 3.9.1 (https://cytoscape.org/).

## Supplemental Movies

Movie S1. Video of PC1 movements in GTP-bound Gα_i1_ Movie S2. Video of PC2 movements in GTP-bound Gα_i1_ Movie S3. Video of PC1 movements in GTP-bound Gα_i2_ Movie S4. Video of PC2 movements in GTP-bound Gα_i2_

## Author Contributions

TJL performed experiments, participated in experimental design, wrote the manuscript. WW performed experiments, participated in experimental design, and edited the manuscript. EM performed experiments, participated in experimental design, and edited the manuscript. SMV performed experiments. NC participated in experimental design.

SA performed experiments, YL performed experiments, CWD participated in experimental design and edited the manuscript, NV participated in experimental design and edited the manuscript, AVS participated in experimental design and edited the manuscript.

## Research Funding

AVS NIH R35GM127303, TJL AHA 826816, NV R01-GM117923 and R01-LM013876

---

**Figure S1.**
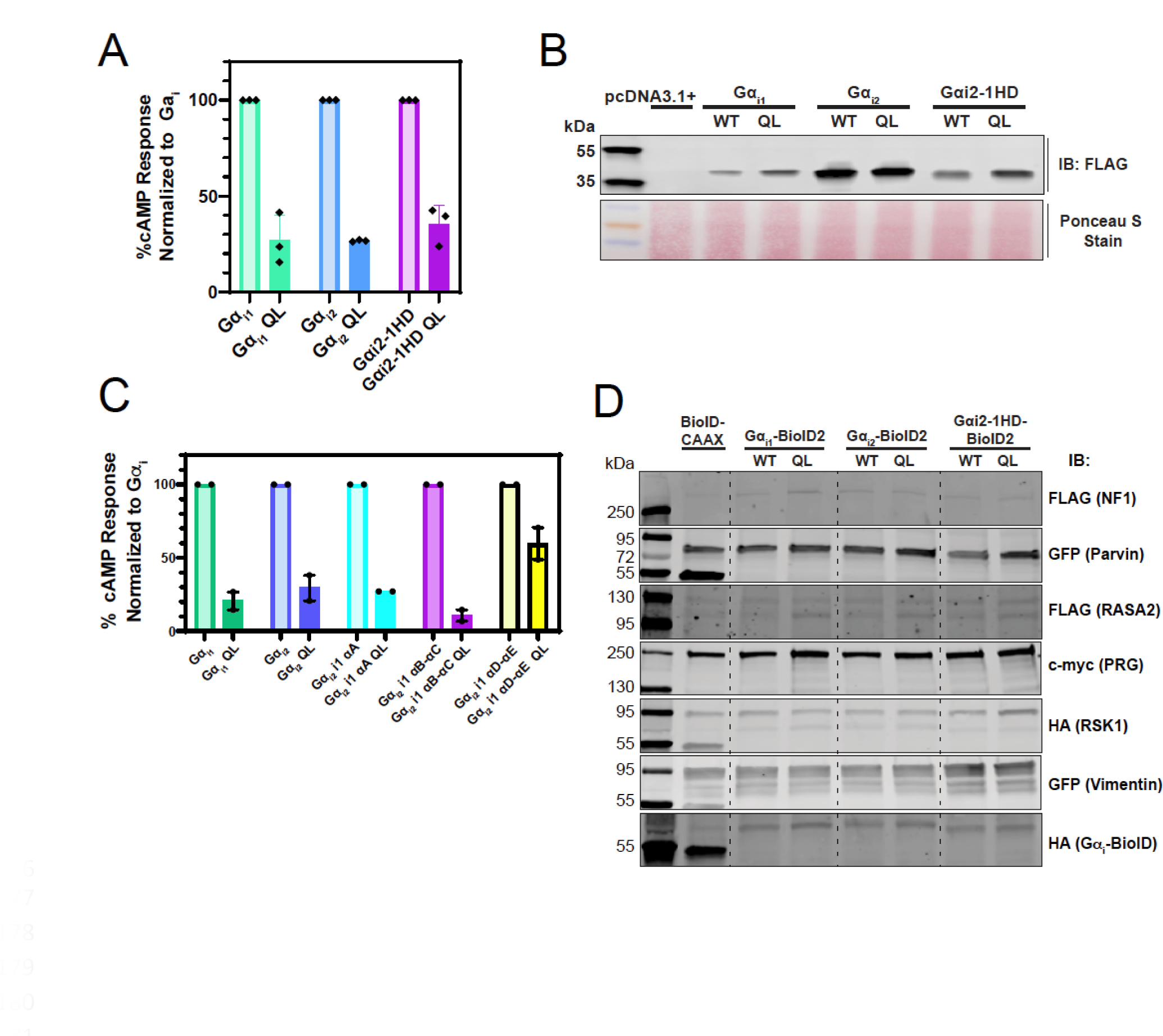
Supporting data for Figure 3. A) cAMP assay measuring the activities of proteins tested in Fig. 3C. B) Western blot showing expression of proteins used in Fig. 3C. C) cAMP assay measuring activities of chimeric proteins used in Fig. 3D. D) Input western blots for BioID2 experiments in Fig. 3E.

**Figure S2.**
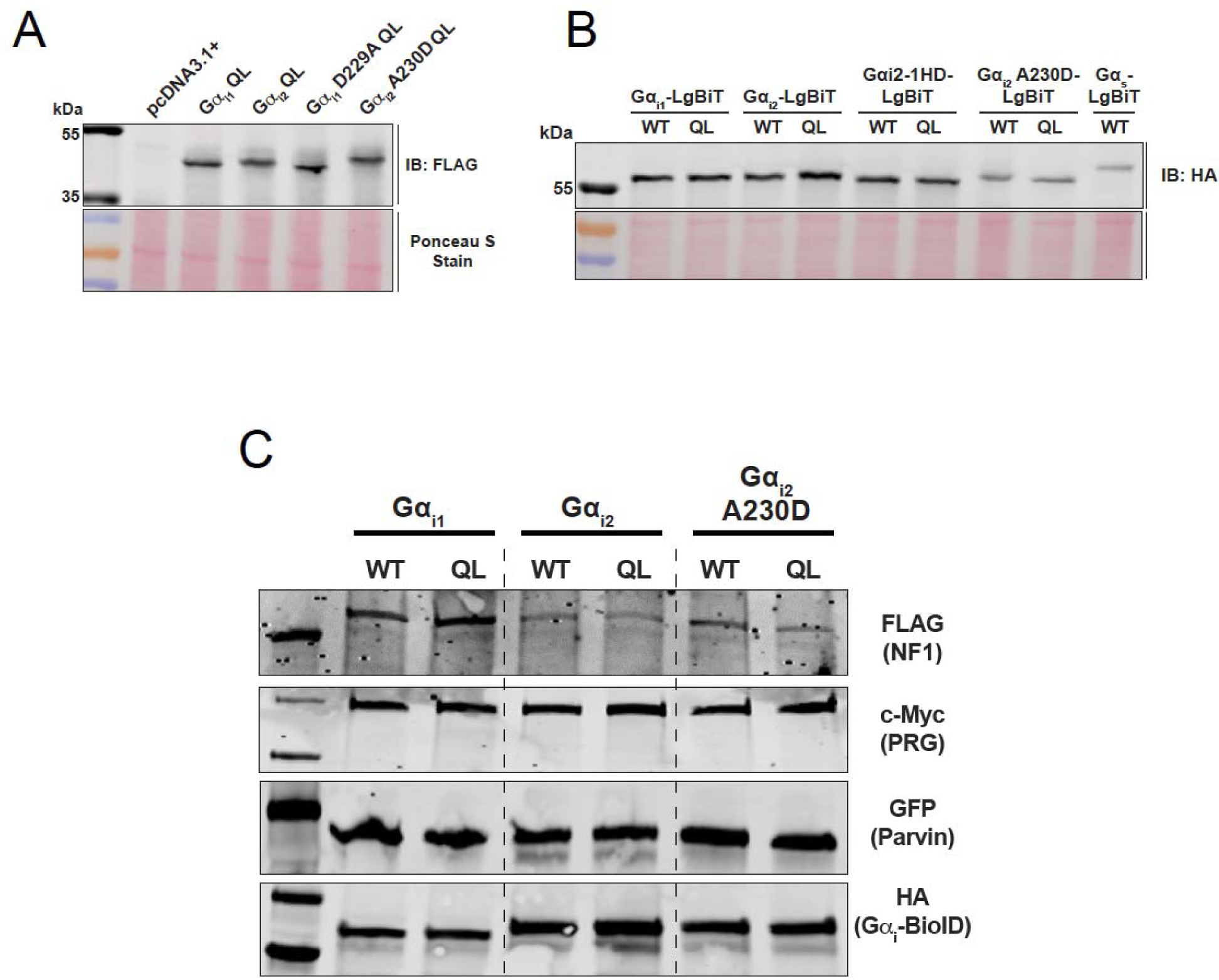
Supporting data for figure 4. A) Western blot for proteins used in Fig. 4B and C. B) Western blot for proteins used in Fig. 4D. C) Input western blots for BioID2 experiments in Fig. 4E.

**Figure S3.**
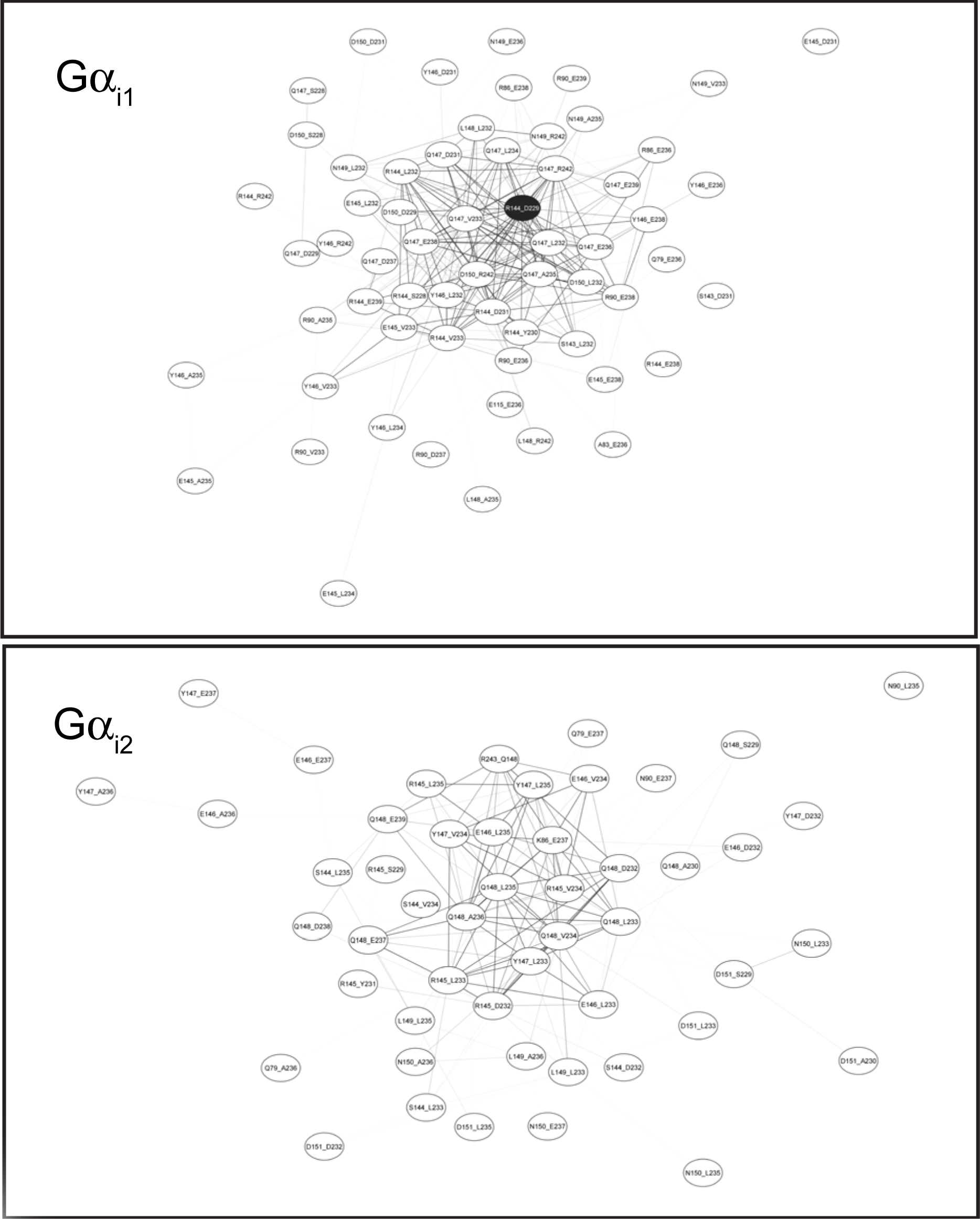
Full Bayesian networks for Gα_i1_ and Gα_i2_ supporting figure 5.

**Figure S4.**
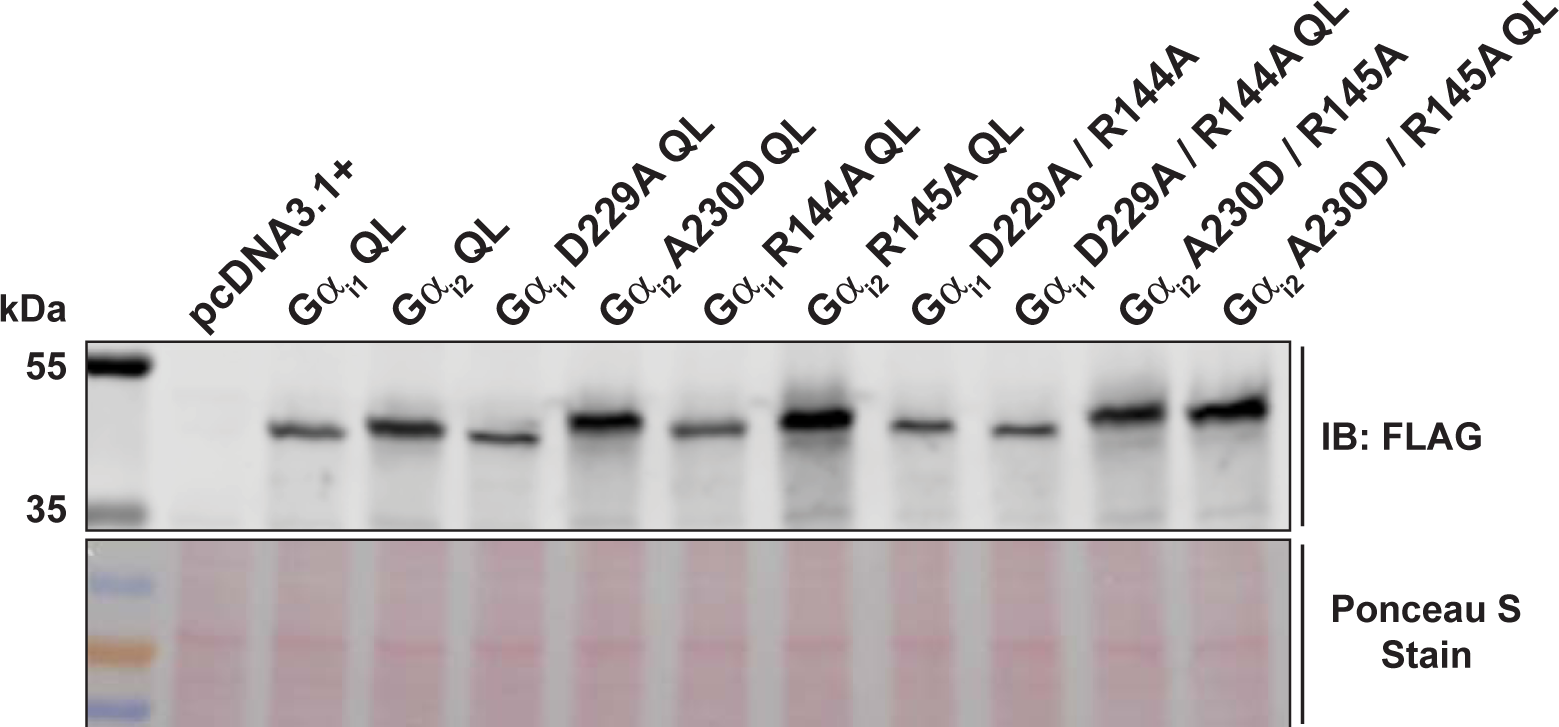
Supporting data for Figure 6. A) Western blot for protein expression for Fig. 6A and B.

**Figure S5.**
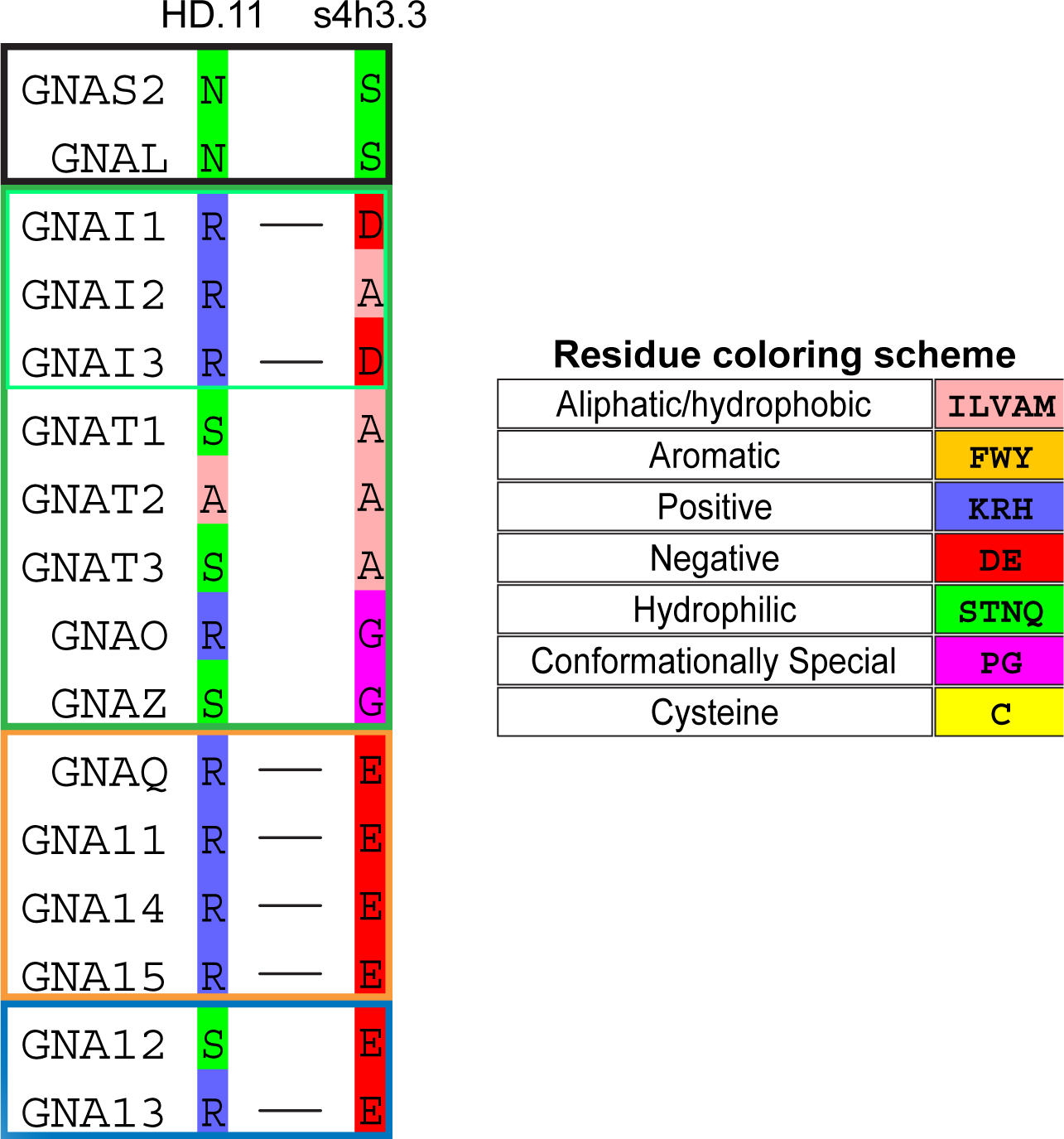
Alignments of all the G protein α subunit families highlighting the presence or absence of ionic lock amino acids HD.11 in the helical domain and s4h3.3 in the RLD.

